# TAK1 Regulates Endothelial Integrity Through Stabilization of Junctions via GSK3β and FoxO1

**DOI:** 10.1101/785907

**Authors:** Sushil C. Regmi, Dheeraj Soni, Dong-Mei Wang, Stephen M. Vogel, Asrar B. Malik, Chinnaswamy Tiruppathi

## Abstract

TLR4 signaling in endothelial cells (ECs) induces vascular injury by disrupting the endothelial junctional barrier. However, it is not known whether TLR4 signaling can also promote endothelial barrier repair after vascular injury. Here we addressed the role of TAK1 activation downstream of TLR4 in the mechanism of vascular integrity. In inducible EC-restricted TAK1 knockout (*TAK1**^iΔEC^***) mice, the endothelial barrier was compromised. Blocking TAK1 activity caused spontaneous loss of the endothelial barrier. Importantly, TAK1 inactivated GSK3β via AKT to prevent β-catenin downregulation. We observed in ECs of *GSK3β^i**ΔEC**^* mice an increase in β-catenin transfer to the nucleus to form a complex with transcription factor FoxO1, thus repressing the expression of the tight junction protein claudin-5 and causing vascular leak. Strikingly, in *TAK1**^iΔEC^*** mice, FoxO1 expression was dramatically increased while expression of AKT was suppressed, and *in vivo* inhibition of FoxO1 prevented sepsis-induced lung vascular leak in *GSK3β^i**ΔEC**^* and *TAK1**^iΔEC^*** mice. Further, EC-restricted deletion of FoxO1 in mice suppressed sepsis-induced lung vascular leak and mortality. Our findings point to the potential of targeting the TAK1-AKT-GSK3β-FoxO1 axis as a therapeutic approach to treat uncontrolled lung vascular leak in sepsis.

## Introduction

Endothelial barrier dysfunction results in protein-rich edema formation and inflammatory cell infiltration that characterize diseases such as acute respiratory distress syndrome (ARDS) (1, 2). In the inflammatory context, endothelial barrier integrity is highly dependent on the processes governing endothelial barrier repair. VE-cadherin (VE-cad) forms Ca^2+^-dependent homophilic *cis* and *trans* dimers at adherens junctions (AJs) that establish cell-to-cell adhesion, which is essential for endothelial barrier integrity (3–5). The cytoplasmic domain of VE-cad interacts with β-catenin (β-cat), which in turn interacts with the actin-associated protein α-catenin (3, 4). The association between β-cat and VE-cad is essential for the formation of endothelial AJs (3, 4). In septic patients and models of sepsis, the lungs are the first organ to fail because of the loss of endothelial barrier function (2). In addition to AJs, the endothelial tight junctions (TJs) also play a critical role in endothelial barrier stability (5–10). Claudin-5 and occludin are the major TJ proteins in endothelial cells (5, 7). Communication between AJs and TJs are thought to be essential for normal endothelial barrier integrity and stability (5, 6). Phosphorylation of AJ protein VE-cad and its associated protein β-cat promotes loss of endothelial barrier integrity (11–16). Similar to AJs, phosphorylation of TJ proteins also causes endothelial barrier disruption (17, 18). Despite both AJs and TJs being important for endothelial integrity, little is known about the signaling mechanisms that regulate endothelial barrier integrity through the control of the distinct set of proteins.

TAK1 is crucial in mediating inflammatory signals activated by cytokines and Toll-like receptor ligands (19–21). TAK1 lies upstream of IκB kinase, p38 MAPK, JNK, and ERK, and causes the activation of NF-κB and MAPK signaling pathways (21, 22). In B cells, TAK1 is required for B cell development and activation of NF-κB and MAPK signaling in response to TLR ligands, and BCR stimuli (23–25); whereas TAK1 in neutrophils negatively regulates NF-κB and p38MAP kinase activation (26). A study using myeloid-specific TAK1 knockout mice showed that TAK1 restricts spontaneous NLRP3 inflammasome activation and thereby prevents cell death in macrophages (27). TAK1 also suppresses RIPK1-dependent melanoma cell death (28) and RIPK3-dependent endothelial necroptosis (29). Global as well as endothelial cell (EC)-specific TAK1 deletion in mice resulted in embryonic lethality (24, 30). Inducible deletion of TAK1 in brain ECs suppressed interleukin 1β (IL-1β)-induced fever and lethargy (31). Deletion of TAK1 in ECs in a tamoxifen-inducible manner in adult mice showed that endothelial TAK1 was essential in preventing TNF-*α*-induced endothelial cell apoptosis (32). Here we investigated the role of TAK1 in regulating endothelial barrier integrity after vascular injury. We observed using inducible EC-restricted TAK1 knockout (*TAK1**^iΔEC^***) mice that endothelial TAK1 was vital for maintaining endothelial barrier integrity and repair of the endothelial barrier after vascular injury. TAK1 functioned by preventing the activation of both GSK3β and FoxO1 via AKT. In *TAK1**^iΔEC^*** mice, we observed a several-fold increase in expression of FoxO1. We also observed using *GSK3β**^iΔEC^*** a defective endothelial barrier secondary to β-cat upregulation. Thus, targeting the TAK1-AKT-GSK3β-FoxO1 axis is a potential therapeutic approach to treat the uncontrolled vascular leak in inflammatory diseases.

## Results

### Endothelial deletion of TAK1 disrupts the endothelial barrier and induces inflammation

Studies were made in tamoxifen-inducible EC-restricted TAK1 knockout (*TAK1^iΔEC^*) mice. *TAK1^iΔEC^*mice were created by breeding loxP-flanked *Map3k7* (*TAK1^fl/fl^*) mice (33) with inducible Cre-ER(T) driven by the 5’endothelial enhancer of stem cell leukemia locus (34). TAK1 was not expressed in ECs of *TAK1^iΔEC^* mice (**Supplemental Fig. 1A-D**). Hematoxylin-and-eosin staining of lung sections from *TAK1^iΔEC^* showed perivascular infiltration of inflammatory cells and hemorrhage as compared with tamoxifen treated *TAK1^fl/fl^* mice (hereinafter referred to as *TAK1^fl/fl^* or WT mice) (Fig. 1A). To determine whether EC-expressed TAK1 affects vascular barrier integrity, we assessed Evans blue albumin (EBA) tracer uptake in lung tissue. EBA uptake increased in *TAK1^iΔEC^* mice compared with *TAK1^fl/fl^*mice (Fig. 1B). Pulmonary transvascular fluid filtration coefficient (*K_f,c_*), a measure of endothelial permeability, increased 8-fold in *TAK1^iΔEC^* lungs compared with WT (Fig. 1C). EC-specific TAK1 deletion also increased mortality in response to LPS induced endotoxemia (Fig. 1D) as well as polymicrobial sepsis induced by cecal ligation and puncture (CLP) as compared to control *TAK1^fl/fl^* mice (Fig. 1E).

**Figure 1.**
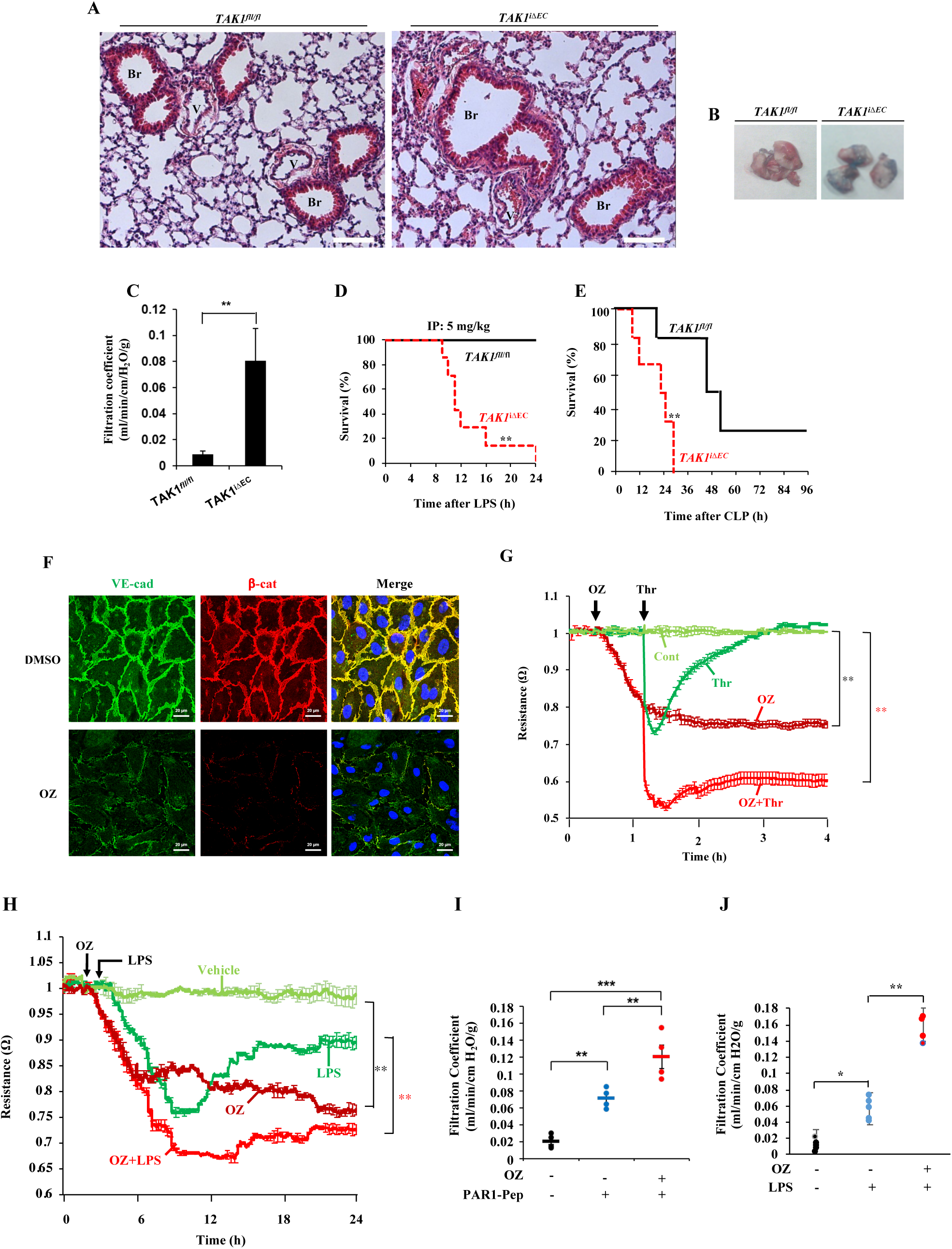
**A-D. EC-restricted TAK1 deletion in mice causes basal lung vascular leak and increases susceptibility to sepsis. A**, Hematoxylin-and-eosin staining of lung sections from *TAK1^fl/fl^* and *TAK1^iΔEC^* mice. scale bars, 100 μm; Br, Bronchi; V, vessel. **B**, *In vivo* lung vascular leak in *TAK1^fl/fl^* and *TAK1^iΔEC^*mice was assessed by measuring Evans blue dye-conjugated albumin (EBA) uptake. Representative photographs of the lungs are shown. **C**, Pulmonary transvascular liquid filtration coefficient in lungs of *TAK1^fl/fl^*and *TAK1^iΔEC^* mice. N = 5 mice per genotype. **p< 0.001, *TAK1^fl/fl^* vs *TAK1^iΔEC^*. **D**, Survival of age- and sex-matched *TAK1^fl/fl^* and *TAK1^iΔEC^*mice after administration of LPS (5 mg/kg, i.p.). N = 10 mice per genotype; **p< 0.001, compared with *TAK1^fl/fl^* (log-rank test). **E**, Survival of age- and sex-matched *TAK1^fl/fl^* and *TAK1^iΔEC^* mice after CLP. N = 10 mice per genotype; *p< 0.01, compared with *TAK1^fl/fl^*(log- rank test). **Figure 1F-J. Inhibition of TAK1 kinase activity causes loss of endothelial barrierfunction F,** HLMVECs grown on coverslips were treated with 1 μM OZ or vehicle (control) for 1 h at 37°C. After this treatment, cells were stained with antibodies specific to VE-cad (green) and β-cat (red), DAPI (blue), and analyzed by confocal microscopy. Results shown are representative of three separate experiments. **G** and **H**, HLMVECs grown to confluence were used to measure real time changes in TER to assess endothelial barrier integrity. Arrows indicate the time at which OZ (1 μM), vehicle (DMSO), thrombin (25 nM) **(G)** or LPS (1μg/ml) (**H**) were added. Values from 3 experiments were presented as mean ± SEM. **p< 0.001, compared with control. **I,** PAR-1 agonist peptide-induced pulmonary transvascular liquid filtration coefficient (*K_f,c_*) in WT mice was measured in the presence and absence of TAK1 inhibitor OZ. Lungs were perfused for 30 min with OZ (1 μM) or vehicle (DMSO) and then PAR-1 agonist peptide was used to induce an increase in *K_f,c_*. N = 5 mice per group. **p< 0.001; ***p< 0.0001. **J**, WT mice were challenged with LPS (5 mg/kg, i.p.) or saline for 6 h and then lungs harvested were used for *K_f,c_* measurement as above with or without OZ (1 μM) pretreatment. N = 5 mice per group. *p< 0.01; **p< 0.001.

To address whether the endothelial barrier disruption in *TAK1^iΔEC^* mice is the result of endothelial death, we performed terminal deoxynucleotidyl transferase dUTP nick end labeling (TUNEL)-positive staining. We did not observe TUNEL-positive cells in lungs of either *TAK1^iΔEC^* or *TAK1^fl/fl^* mice (**Supplemental Figure 1E**). Knockdown of TAK1 in human lung microvascular ECs (ECs) using small interfering RNA (siRNA) also did not induce apoptosis (**Supplemental Figure 1F)** indicating that TAK1 deficiency did not injure the endothelium due to apoptosis.

### Inhibition of TAK1 activity disrupts endothelial barrier

Since *TAK1^iΔEC^* mice showed increased endothelial permeability reflecting endothelial injury, we next determined whether TAK1 kinase activity was responsible for endothelial barrier disruption. On inhibiting TAK1 kinase activity with 5Z-7-oxozeaenol (OZ) (1 μM, 1 h), we observed marked reductions in the expression of VE-cad and β-catenin at AJs in ECs (Fig. 1F). Measurement of trans-endothelial resistance (TER) showed that OZ (1 μM) itself caused a time dependent decrease in TER (Fig. 1G, 1H) indicating disassembly of endothelial AJs. We observed that the protease-activated receptor-1 (PAR-1) agonist thrombin (generated in vasculature during sepsis) produced a transient increase in permeability whereas LPS produced a sustained increase in permeability as reflected by the relative decreases in TER; however, both responses were more sustained and greater with OZ treatment as compared with controls (Fig. 1G, 1H). Next, to address the *in vivo* effect of TAK1 kinase inhibition, we measured PAR-1-induced changes in *K_f,c_* in mouse lungs and observed that the PAR-1 agonist peptide (TFLLRNPNDK-NH_2_) induced a 3-fold increase in *K_f,c_* over the basal value in WT mouse lungs (Fig. 1I), whereas in mouse lungs perfused with OZ (1 μM) for 30 min the PAR-1 agonist significantly augmented the increase in *K_f,c_* (Fig. 1I). WT mice, challenged with a low dose of LPS (5 mg/kg, i.p.,) for 6 h and then exposed to OZ as above, also showed a similar augmentation of increase in endothelial permeability (Fig. 1J). These findings show that TAK1 kinase activity functions to suppress endothelial barrier injury in response to LPS and thrombin.

### TAK1 inhibition of GSK3β prevents β-catenin degradation and maintains endothelial permeability

To examine mechanisms of TAK1 induced endothelial barrier protection, we focused on the expression of adherens junction protein β-catenin (β-cat) in TAK1 deficient ECs. Here we used ECs from *TAK1^fl/fl.Cre+^* mice that were treated with tamoxifen (2 μM) for 72 h to delete TAK1 (Fig. 2A), which reduced β-cat expression by 80% as compared with control ECs (Fig. 2B). β-cat contains several consensus phosphorylation sites (amino acid positions: S29, S33, S37, T4, S45) for GSK3β (35). GSK3β phosphorylation of β-cat promotes β-cat ubiquitination and degradation via the proteasomal pathway (35, and inhibition of GSK3β has been linked to enhanced endothelial barrier function due to increased β-cat expression at AJs (37). Thus, we addressed whether TAK1 inhibited GSK3β and thereby stabilized β-cat at AJs to promote barrier function. We first determined the effects of OZ, the inhibitor of TAK1 activity, on the phosphorylation of GSK3β on S9 to inactivate GSK3β in ECs (Fig. 2C, 2D). OZ prevented basal as well as thrombin- and LPS-induced phosphorylation of GSK3β at S9 (Fig. 2C, 2D). Furthermore, knockdown of TAK1 also prevented phosphorylation of GSK3β at S9 in ECs (Fig. 2E), suggesting that it would lead to increased phosphorylation of β-cat, increased β-cat degradation and hence decreased β-cat expression at AJs. To address whether this was the case, we determined the phosphorylation of S33, S37 and T41 on β-cat in control or OZ treated ECs (Fig. 2F, 2G). We observed that thrombin and LPS both induced phosphorylation of β-cat in a time dependent manner in ECs (Fig. 2F, 2G) and that OZ increased β-cat phosphorylation at these sites (Fig. 2F, 2G). To address next whether the proteasomal pathway was responsible for β-cat degradation, we determined the effects of TAK1 inhibition on β-cat ubiquitination. ECs were incubated for 2 h with the proteasomal inhibitor MG-132 and then challenged either with thrombin or LPS. We observed that basal as well as thrombin- and LPS-induced ubiquitination of β-cat was increased in OZ-treated ECs (Fig. 2H, 2I). Since OZ treatment disrupted the endothelial barrier (Fig. 1F-1J), we determined the effect of the GSK3β inhibitor (SB216763) on OZ induced endothelial barrier disruption. Pretreatment with the GSK3β inhibitor prevented the OZ-induced loss of VE-cad and β-cat at AJs (Fig. 2J) as well as OZ-plus thrombin-induced decreases in TER (Fig. 2K). These results show that TAK1 inhibition of GSK3β prevents β-cat degradation and hence promotes β-cat localization at AJs and maintains endothelial permeability.

**Figure 2.**
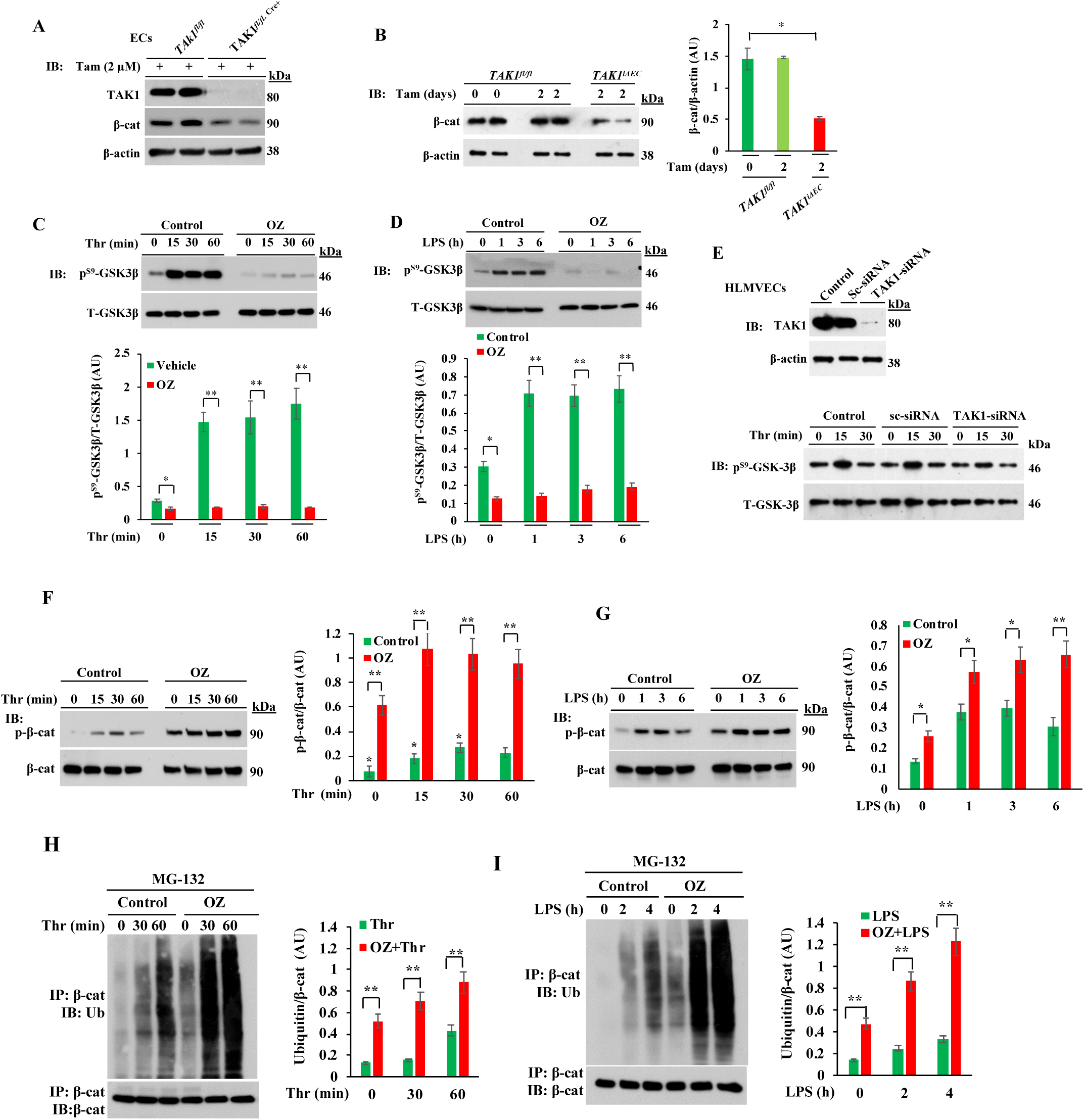
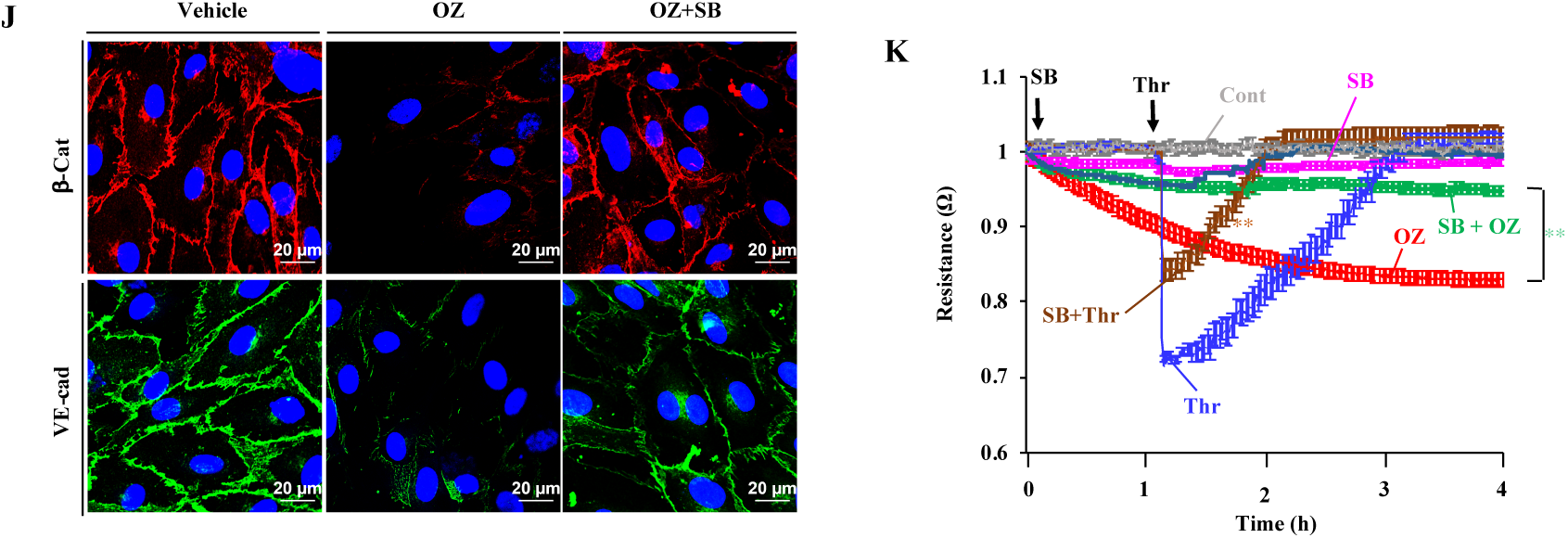
TAK1 stabilizes β-catenin at endothelial AJs by inhibiting GSK3β. **A**, LECs from *TAK1^fl/fl^* and *TAK1^fl/fl.Cre+^* mice in culture were treated with tamoxifen (Tam) for 72 h and then used for immunoblot (IB) analysis. Results shown are representative of three separate experiments. **B**, *TAK1^fl/fl^*and *TAK1^fl/fl.Cre+^* mice were injected i.p. with tamoxifen (1 mg/mouse/day) for 2 days and at 7^th^ day, lungs harvested were used for IB analysis. Results shown are representative blots of lung extracts from 6 mice per genotype. *p<0.01, compared with *TAK1^fl/fl^*mice. **C** and **D**, HLMVECs were treated with or without OZ (1 μM) for 1 h and then exposed to either thrombin (25 nM) or LPS (1μg/ml). Cell lysates were used for IB analysis to determine GSK3β phosphorylation on S-9. *p< 0.05; **p< 0.001, compared with vehicle used. **E**, HLMVECs transfected with either control-siRNA (Sc-siRNA) or TAK1-siRNA were used for determination of thrombin-induced phosphorylation of GSK3β on S-9. *Top panel* shows that TAK1 expression was suppressed in TAK1-siRNA treated cells. *Bottom panel* shows that TAK1 knockdown suppresses thrombin-induced S-9 phosphorylation of GSK3β. Results shown are representative of 3 experiments. **F** and **G**, HLMVECs were treated with or without OZ (1 μM) for 1 h and then thrombin- or LPS-induced phosphorylation of β-cat on S33, S37, and T41 was measured. Results shown are representative of three separate experiments. Quantified data are presented as the mean ± SEM ratio of phosphorylated:total protein after OZ treatment *vs* control. *p< 0.01; **p< 0.001. **H** and **I**, HLMVECs incubated with proteasomal inhibitor MG132 (10 μM) for 2 h followed by 1 μM OZ for 1 h were exposed to either thrombin (25 nM) (**H**) or LPS (1 μg/ml) (**I**) for indicated time periods. Then cell lysates were immunoprecipitated with anti-β-cat Ab and blotted with anti-pan ubiquitin Ab. Quantified data from 3 experiments are presented as the mean ± SEM of ubiquitin: β-cat ratio in OZ treated vs control. **p< 0.001. **J**, HLMVECs were treated with vehicle, OZ (1 μM), or OZ (1 μM) plus SB216763 (1 μM) for 1 h and then cells stained with antibodies specific to β-cat (red), VE-cad (green), and nuclear DAPI (blue). Then cells were analyzed by confocal microscopy. **K,** HLMVECs grown to confluence treated with vehicle (control), 1 μM OZ, or 1 μM OZ plus 1 μM SB (SB216763) were used to measure real-time changes in TER to assess endothelial barrier integrity. Arrows indicate the time at which inhibitors or thrombin (25 nM) were added. **p< 0.001, OZ vs SB + OZ, or thrombin alone vs SB + thrombin.

### Endothelial-specific GSK3β deletion-dependent nuclear localization of β-catenin and FoxO1 increases endothelial permeability

Next, to examine the *in vivo* role of GSK3β in ECs and to determine its role in regulating endothelial permeability, we generated tamoxifen-inducible EC-restricted GSK3β knockout (*GSK3β**^iΔEC^***) mice (**Supplemental Figure 2A-D).** We observed augmented β-cat expression in ECs from *GSK3β^iΔEC^* mice compared to WT littermates (*GSK3β^fl/fl^*) (Fig. 3A), indicating that phosphorylation of β-cat and its degradation are reduced in the absence of GSK3β in ECs. We determined the effects of deletion of *GSK3β* on vascular permeability using *GSK3β^iΔEC^*. Basal as well as PAR-1- and LPS-induced increases in permeability were augmented in *GSK3β^iΔEC^* mice compared with WT mice (Fig. 3B, 3C). In addition, polymicrobial sepsis induced by CLP produced 100% mortality in *GSK3β^iΔEC^* mice (Fig. 3D), whereas CLP induced only 20% mortality in *GSK3β^fl/fl^* mice (Fig. 3D). LPS dose of 10 mg/kg i.p. produced 100% mortality in *GSK3β^iΔEC^*mice (Fig. 3E) but the same challenge produced no mortality in *GSK3β^fl/fll^* mice (Fig. 3E).

**Figure 3.**
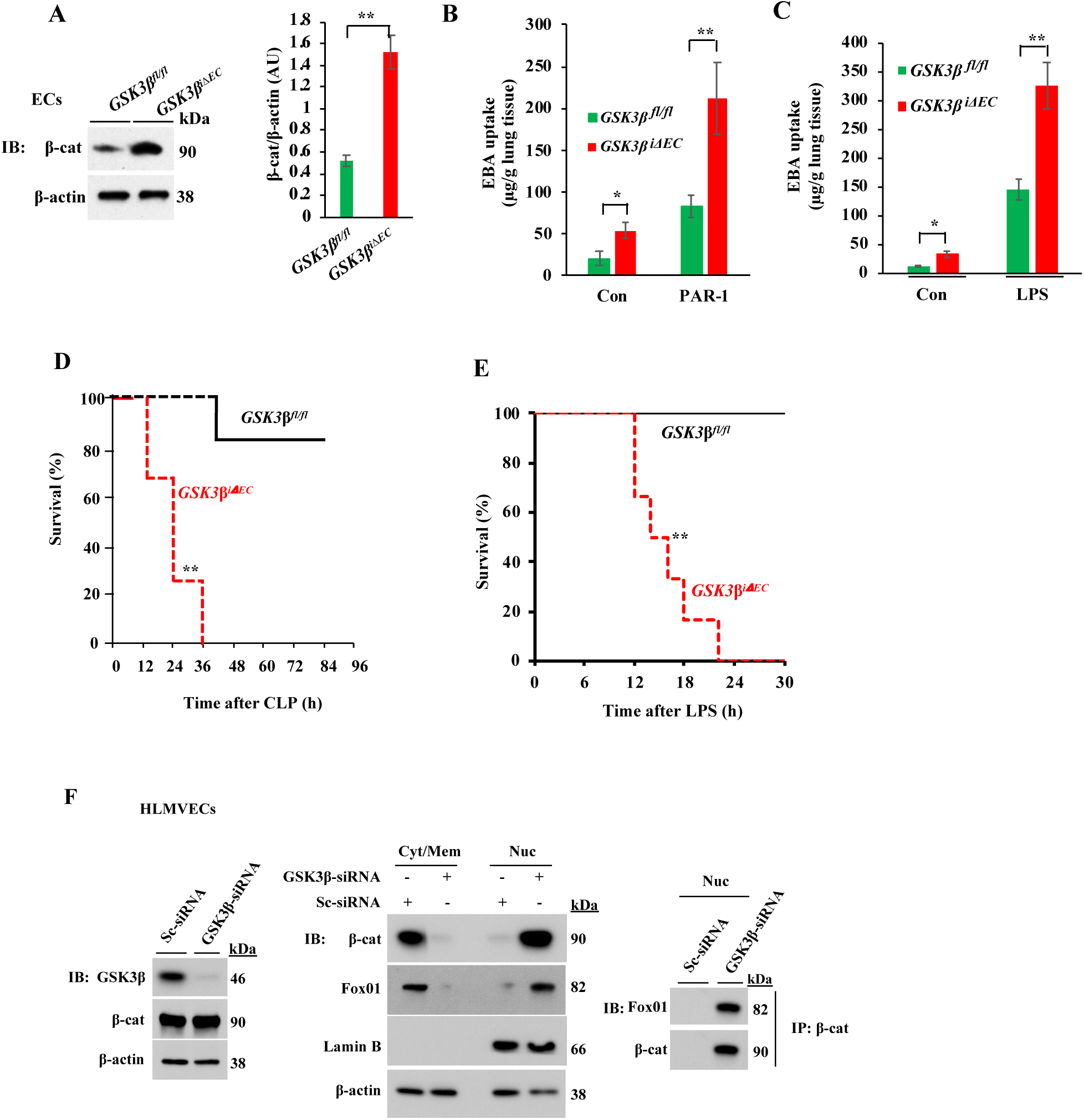
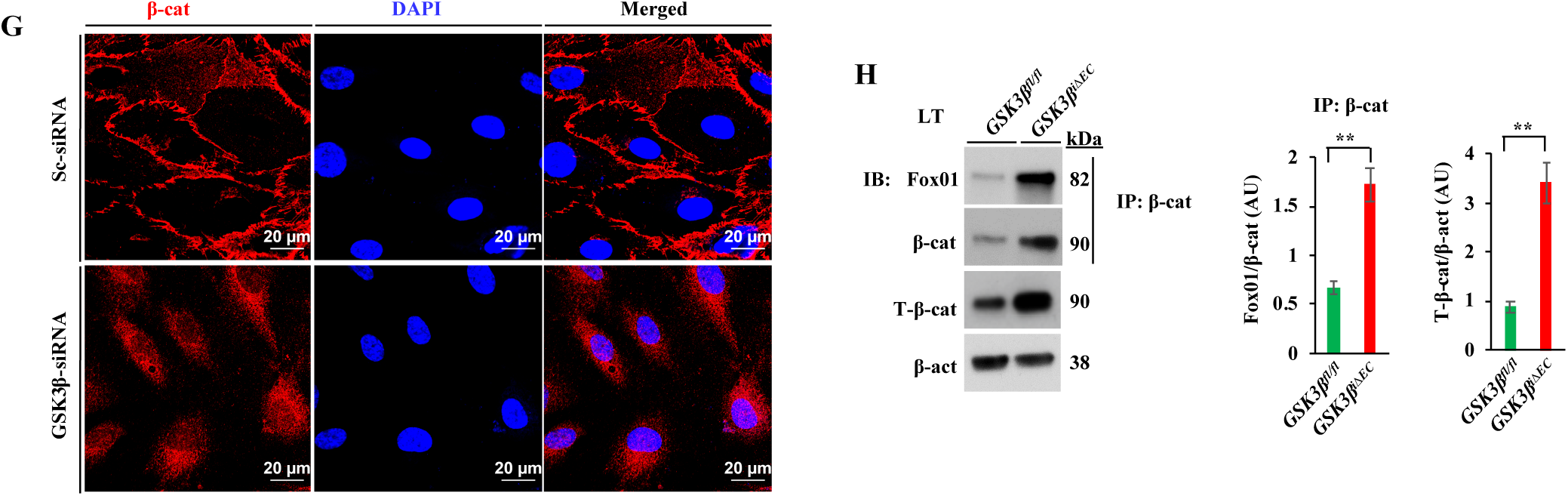
GSK3β deficiency in ECs promotes loss of endothelial barrier function via activating transcriptional repressor complex β-catenin/FoxO1. **A**, Lung ECs from GSK3β*^fl/fl^* and GSK3β*^iΔEC^* mice were used for IB analysis to determine β-cat expression. N = 5 mice per genotype; **p<0.001. **B**, PAR-1 agonist peptide-induced lung vascular leak in GSK3β*^fl/fl^* and GSK3β*^iΔEC^* mice was assessed by measuring EBA uptake. N = 5 mice per genotype. *p< 0.01; **p< 0.001; GSK3β*^fl/fl^* vs GSKβ*^iΔEC^* mice. **C**, GSK3β*^fl/fl^* and GSK3β*^iΔEC^* mice were challenged with LPS (5 mg/kg, i.p.) or saline for 6 h and then *in vivo* EBA uptake in lungs was measured. *p< 0.01; **p< 0.001; GSK3β*^fl/fl^* vs GSKβ*^iΔEC^* mice. **D**, Survival of age- and weight-matched *GSK-3*β *^fl/fl^* and *GSK3*β *^iΔEC^* mice after CLP. n = 10 mice per genotype. **p< 0.001 (log-rank test). **E**, Survival of age- and weight-matched *GSK3β ^fl/fl^* and *GSK3β ^iΔEC^* mice after administration of LPS (10 mg/kg, i.p.). n = 10 per genotype. **p< 0.001 (log-rank test). **F**, HLMVECs were transfected with 100 nM scrambled (Sc)-siRNA or GSK3β-siRNA. At 48 h after transfection, cells were used for IB analysis to determine GSK3β expression (left panel) or used to prepare nuclear and cytosolic/membrane fractions to determine β-cat and FoxO1 expression in cytosol/membrane and nucleus by IB (*middle panel*). Lamin B, nuclear marker. Nuclear fractions from Sc-siRNA and GSK3β-siRNA transfected cells were lysed, immunoprecipitated with anti-β-cat pAb and blotted with anti-FoxO1 mAb (*right panel*). **G**, HLMVECs transfected with Sc-siRNA or GSK3β-siRNA were stained with anti-β-cat Ab (red) and analyzed by confocal microscopy. DAPI, nuclear marker (blue). **H**, Lung lysates from *GSK3β ^fl/fl^* and *GSK3β ^iΔEC^* mice were immunoprecipitated with anti-β-cat pAb and blotted with anti-FoxO1 mAb. N = 4 mice per genotype; **p< 0.001, *GSK3β ^fl/fl^* vs *GSK3β ^iΔEC^*.

To address possible alteration in overall β-cat expression in *GSK3β^iΔEC^* mice, we measured the subcellular distribution of β-cat. GSK3β depletion markedly reduced membrane associated β-cat expression at AJs whereas β-cat was diffusely localized in nuclei and cytoplasm (Fig. 3F, 3G). Immunoblotting experiments showed that depletion of GSK3β induced a shift in the localization of β-cat from the cytosol to nuclear fraction as compared with control cells (Fig. 3F). This was also seen with the transcription factor FoxO1 (Fig. 3F) consistent with previous evidence (5, 6). In the absence of GSK3β in ECs, anti-β-cat antibody immunoprecipitated both β-cat and FoxO1 as compared to ECs from WT mice (Fig. 3H).

### FoxO1 inhibition restores the endothelial barrier in *GSK3β^iΔEC^*mice

To determine whether FoxO1/β-cat interaction in the nucleus is responsible for the loss of the endothelial barrier in *GSK3β^iΔEC^*mice, we analyzed the lung lysates from *GSK3β^fl/fl^* and *GSK3β^iΔEC^*mice. Under basal conditions, FoxO1 expression was increased in *GSK3β^iΔEC^* mice (Fig. 4A); in contrast basal expression of claudin-5, occludin, and VE-cad were strikingly low in *GSK3β^iΔEC^* mice compared with basal levels in WT mice (Fig. 4C, 4D). However, β-cat expression was increased in *GSK3β^iΔEC^* mice compared with WT mice (Fig. 4D). Next, we studied the effect of FoxO1 inhibitor (AS1842856; it binds to the active site on FoxO1 to inhibit transcriptional activity of FoxO1 [38]) on endothelial barrier stability *in vivo*. Here, we treated WT and *GSK3β^iΔEC^* mice with the FoxO1 inhibitor and then challenged them with LPS to assess endothelial barrier stability. In WT mice, LPS markedly reduced the expression of claudin-5, VE-cad, and β-cat (Fig. 4C, 4D). Interestingly, FoxO1 inhibitor treatment prevented LPS-induced loss of claudin-5, VE-cad, and β-cat expression in the lung (Fig. 4C, 4D) as well as lung vascular leak as assessed by EBA uptake in WT mice (Fig. 4E). Similarly, we observed that in *GSK3β^iΔEC^* mice, FoxO1 inhibitor treatment restored the expression of claudin-5, occludin, VE-cad, and β-cat to normal levels (Fig. 4C, 4D) and prevented LPS-induced lung vascular leak (Fig. 4E). Thus, the mechanism of increased permeability seen following deletion of GSK3β and nuclear translocation of β-cat and β-cat/FoxO1 interaction involved reduced expression of adherens and tight junction proteins.

**Fig. 4.**
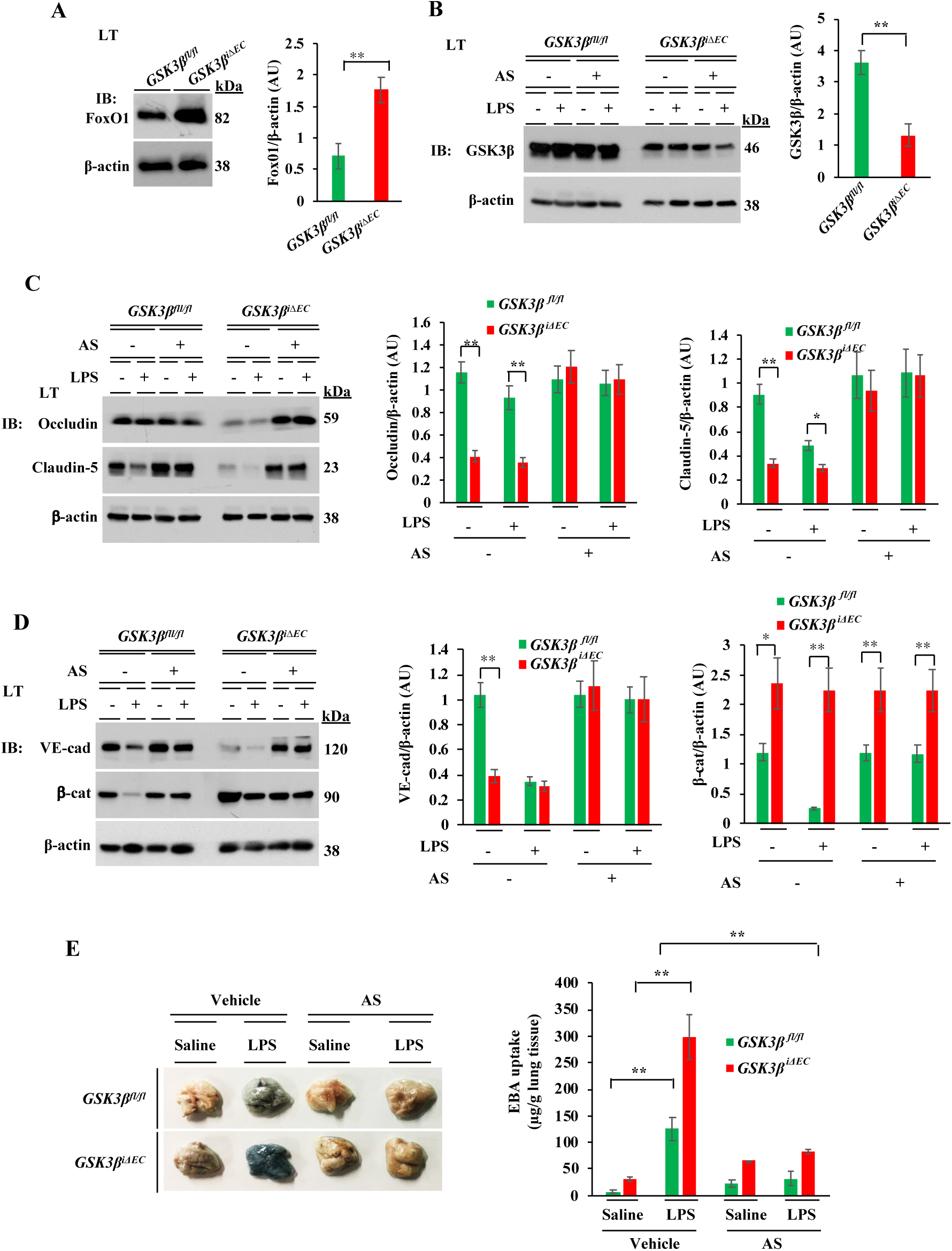
FoxO1 inhibition restores the endothelial barrier in *GSK3β^iΔEC^*mice. **A**, Lung lysates from *GSK3β ^fl/fl^* and *GSK3β ^iΔEC^* mice were used for IB analysis to determine FoxO1 expression. N = 5 mice per genotype; **p< 0.001, *GSK3β ^fl/fl^ vs GSK3β ^iΔEC^* mice. **B-D,** *GSK3β ^fl/fl^* and *GSK3β ^iΔEC^* mice injected i.p. either vehicle (DMSO) or FoxO1 inhibitor AS1842856 (AS; 5 mg/kg, i.p. one injection per day for 4 consecutive days) were challenged with i.p. LPS (5 mg/kg) or saline for 6 h. Lungs harvested were used for IB analysis to determine GSK3*β* (**B**), occludin and claudin-5 (**C**), and VE-cad, and β-cat (**D**). N = 5 mice/genotype/group. *p<0.05; **p< 0.001; *GSK3β ^fl/fl^ vs GSK3β ^iΔEC^* mice. **E,** *GSK3β ^fl/fl^* and *GSK3β ^iΔEC^* mice injected i.p. with either vehicle (DMSO) or FoxO1 inhibitor (AS) were challenged with i.p. LPS (5 mg/kg) or saline for 6 h and used to assess lung vascular leak by measuring EBA uptake. Representative lung images are shown in *left panel*. Quantified results are shown in *right panel.* N = 5 mice/genotype/group. **p<0.001, basal *vs* LPS in *GSK3β ^fl/fl^* mice; LPS challenge, *GSK3β ^fl/fl^ vs GSK3β ^iΔEC^* or LPS challenge vs AS plus LPS challenge.

### TAK1 activates AKT to suppress the function of GSK3β

Since we observed that chemical inhibition of TAK1 kinase blocked thrombin- or LPS-induced phosphorylation of GSK3β at S9, we addressed the possibility that TAK1 inactivates GSK3β through AKT, which is known to phosphorylate GSK3β at S9 to inactivate GSK3β (36). In support of this concept, OZ treatment blocked thrombin- or LPS-induced phosphorylation of AKT (Fig. 5A, 5B). Further, thrombin or LPS stimulation caused interaction between TAK1 and AKT in ECs (Fig. 5C). In addition, an AKT inhibitor prevented thrombin- or LPS-induced S-9 phosphorylation of GSK3β (Fig. 5D, 5E). Also, we observed that the AKT inhibitor blocked thrombin- or LPS-induced phosphorylation of AKT in ECs (Fig. 5F, 5G). In other experiments, we measured the effect of the AKT inhibitor on endothelial barrier function by measuring TER. We observed that AKT inhibition augmented basal as well as thrombin- or LPS-induced decrease in TER (Fig. 5H, 5I). Taken together, these results suggest that TAK1 activates AKT which in turn inactivates GSK3β in ECs.

**Figure 5.**
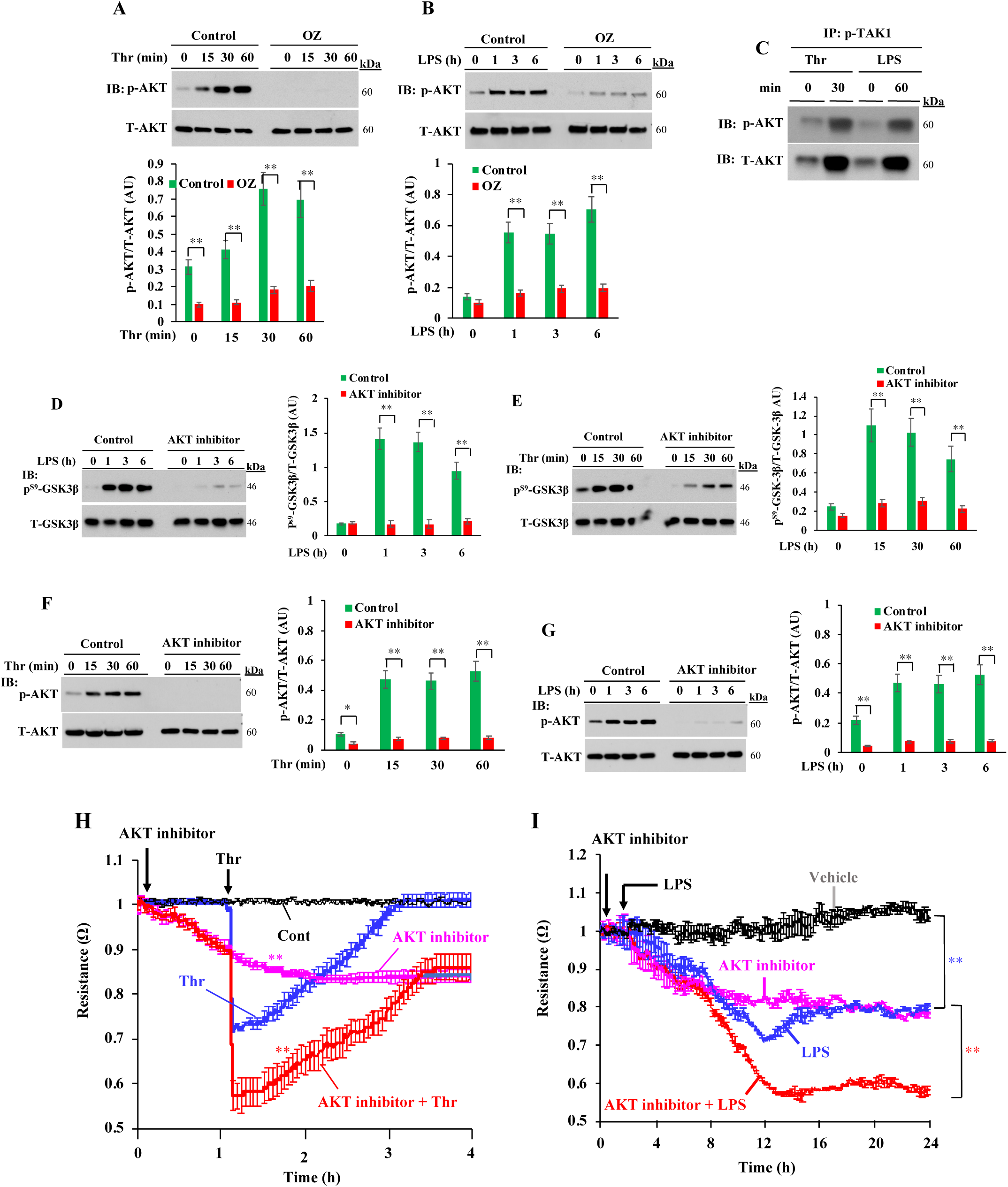
TAK1 inhibits GSK3β through AKT activation. **A-C**, HLMVECs treated with either vehicle (control) or 1 μM OZ for 1 h and then challenged with thrombin (**A**) or LPS (**B**) were used for IB analysis. **p< 0.001, compared with controls. **C**, HLMVECs treated with thrombin or LPS were immunoprecipitated with anti-phospho-TAK1 Ab and blotted with anti-phospho-AKT Ab or anti-AKT Ab. Results shown are representative of 3 separate experiments. **D** and **E**, HLMVECs treated either with vehicle (control) or 1μM AKT inhibitor for 1 h and then challenged with thrombin (**D**) or LPS (**E**) were used for IB analysis to determine S-9 phosphorylation of GSK3*β*. Results shown are representative of 3 separate experiments. **p< 0.001, compared with controls. **F** and **G**, HLMVECs treated either with vehicle (control) or 1 μM AKT inhibitor for 1 h and then challenged with thrombin (**F**) or LPS (**G**) were used for IB analysis to determine phosphorylation of AKT. *p<0.01; **p< 0.001, compared with controls. **H** and **I**, HLMVECs treated with vehicle (control) or 1 μM AKT inhibitor and then thrombin (25 nM) or LPS to induce real-time changes in TER was measured to assess endothelial AJ integrity. Arrows indicate the time at which inhibitors or thrombin (25 nM) or LPS (1 μg/ml) were added. **p< 0.001, AKT inhibitor vs controls, thrombin vs AKT inhibitor plus thrombin, or LPS vs AKT inhibitor plus LPS.

### TAK1 restricts FoxO1 function in ECs through AKT activation

Since AKT-mediated phosphorylation of FoxO1 inhibits FoxO1 DNA binding activity and expels FoxO1 the from nucleus (39–41), we investigated whether TAK1 occludes the function of FoxO1 via AKT. We observed that TAK1 inhibitor OZ blocked both thrombin- and LPS-induced phosphorylation of FoxO1 at T-24 and S-256 in ECs (Fig. 6A, 6B). As a positive control, we showed that a FoxO1 inhibitor (AS1842856 [AS]) also blocked thrombin- or LPS-induced phosphorylation of FoxO1 in ECs (Fig. 6A, 6B). In addition, FoxO1 inhibition largely prevented the spontaneous loss of AJs in ECs (Fig. 6C) due to TAK1 inhibition. Moreover, concurrent inhibition of both GSK3β and FoxO1, completely reversed the OZ induced effect on endothelial AJs (Fig. 6C) as well as the LPS effect on endothelial barrier integrity as assessed by measuring TER (Fig. 6D, 6E). These findings support the proposal that TAK1 controls the function of GSK3β and FoxO1 in ECs.

**Figure 6.**
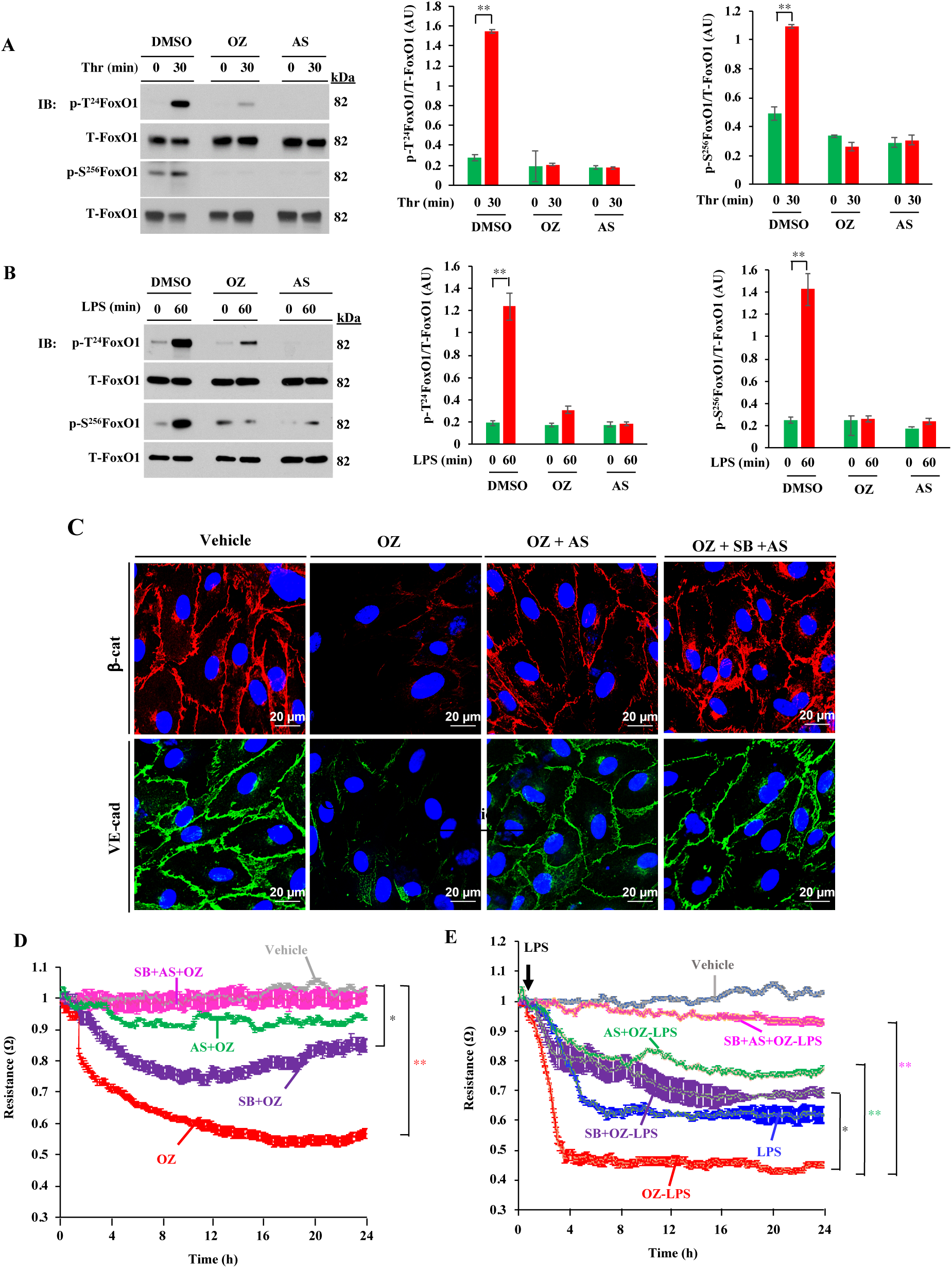
TAK1-mediated FoxO1 inhibition stabilizes endothelial barrier. **A** and **B**, HLMVECs treated with vehicle (control), 1 μM OZ, or 100 nM AS (FoxO1 inhibitor) for 1 h and then challenged with thrombin (**A**) or LPS (**B**) were used for IB analysis to determine phosphorylation of FoxO1 at T-24 and S-256. Results shown are representative of 3 separate experiments. **p< 0.001, compared with unstimulated cells. **C**, HLMVECs were treated with vehicle (control), 1 μM OZ, 1μM OZ + 100 nM AS, or 1 μM OZ + 1 μM SB + 100 nM AS for 1 h, and then cells were stained with antibodies specific to β-cat (red), VE-cad (green) and DAPI (blue), and analyzed by confocal microscopy. Results shown are representative of 3 experiments. **D** and **E**, Endothelial barrier integrity was assessed by measuring TER. HLMVECs grown to confluence were used to measure real-time changes in TER in the presence of vehicle (control), 1 μM OZ, 1 μM OZ + 100 nM AS, or 1 μM OZ + 1 μM SB + 100 nM AS. In **E**, cells treated with inhibitors as in **D** were challenged with LPS (1μg/ml). *p<0.05, control vs SB+OZ; OZ-LPS vs SB+OZ-LPS; **p<0.001, control vs OZ, OZ-LPS vs SB+OZ-LPS, or OZ-LPS vs SB+AS+OZ-LPS.

To define the *in vivo* role of EC-expressed TAK1 in regulating FoxO1, we analyzed lung lysates from *TAK1^fl/fl^* and *TAK1^iΔEC^* mice. Remarkably, we observed a several-fold increase in expression of FoxO1 (Fig. 7A) while the expression of AKT was suppressed in lungs of *TAK1^iΔEC^* mice (Fig. 7A). Since FoxO1 represses the transcription of claudin-5 by associating with β-cat (6), we observed that claudin-5 expression was markedly reduced along with that of occludin, VE-cad, and β-cat in *TAK1^iΔEC^* mice compared with *TAK1^fl/fl^* mice (Fig. 7B, 7C). LPS challenge caused downregulation of TJ and AJ proteins in both genotypes (Fig. 7B, 7C), however, the effect was more dramatic in *TAK1^iΔEC^* mice (Fig. 7B, 7C). Next, we pretreated both genotypes with a FoxO1 inhibitor and then challenged with LPS and observed that FoxO1 inhibition prevented LPS-induced downregulation of TJ and AJ proteins (Fig. 7B, 7C) as well as lung vascular leak in both genotypes (Fig. 7D). To determine whether FoxO1 could be a potential therapeutic target to treat sepsis, we investigated the effect of FoxO1 inhibition on the survival of *TAK1^fl/fl^* and *TAK1^iΔEC^*mice after CLP. We administered the FoxO1 inhibitor or saline i.p. 6 h after CLP and monitored survival of both genotypes. In saline-treated *TAK1^fl/fl^* mice, CLP produced 80% mortality in 4 days whereas in *TAK1^iΔEC^*mice, CLP produced 100% mortality within 30 h (Fig. 7E). Importantly, we observed that FoxO1 inhibitor treatment markedly increased the survival of both *TAK1^fl/fl^* and *TAK1^iΔEC^* mice after CLP (Fig. 7E).

**Figure 7.**
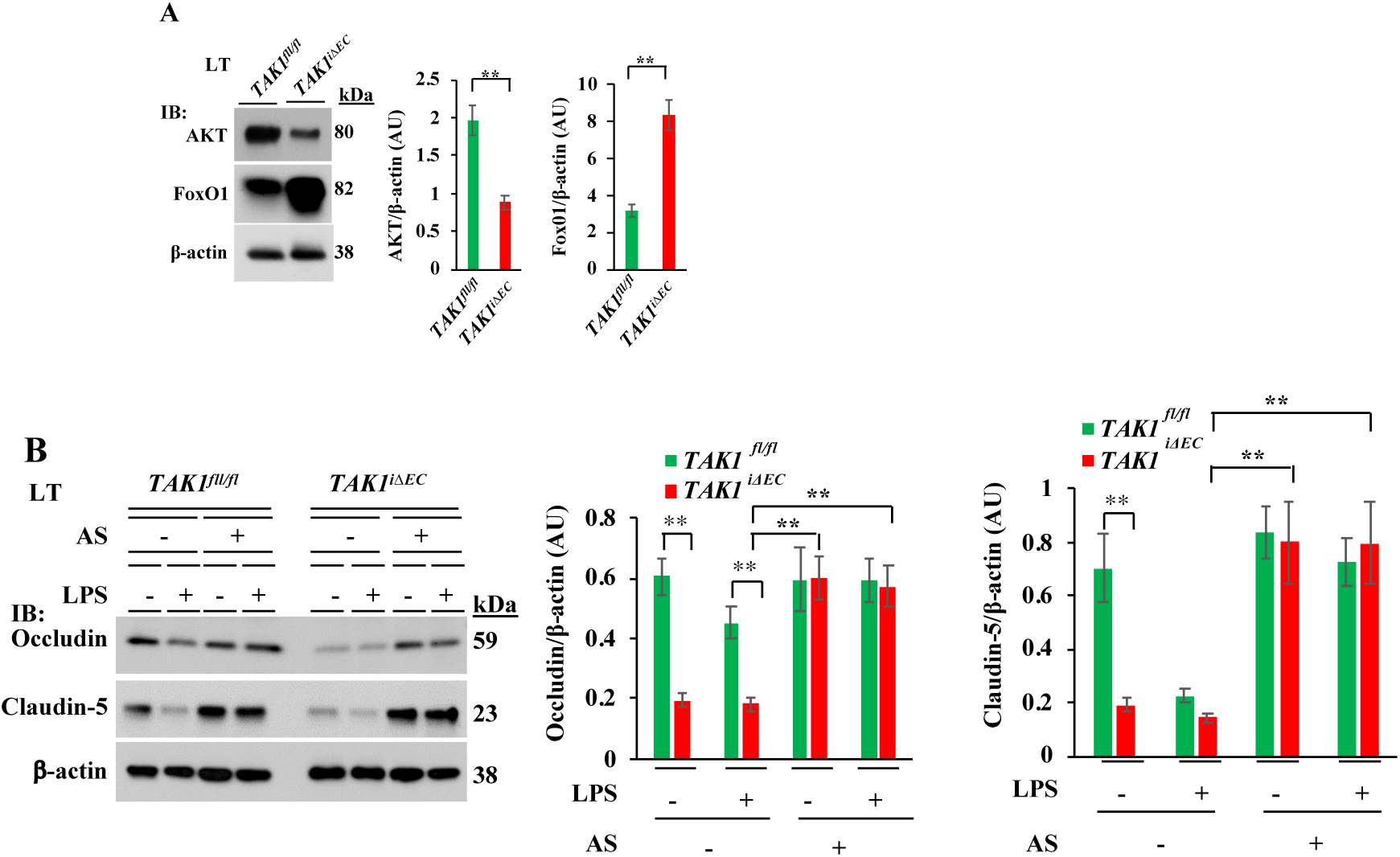
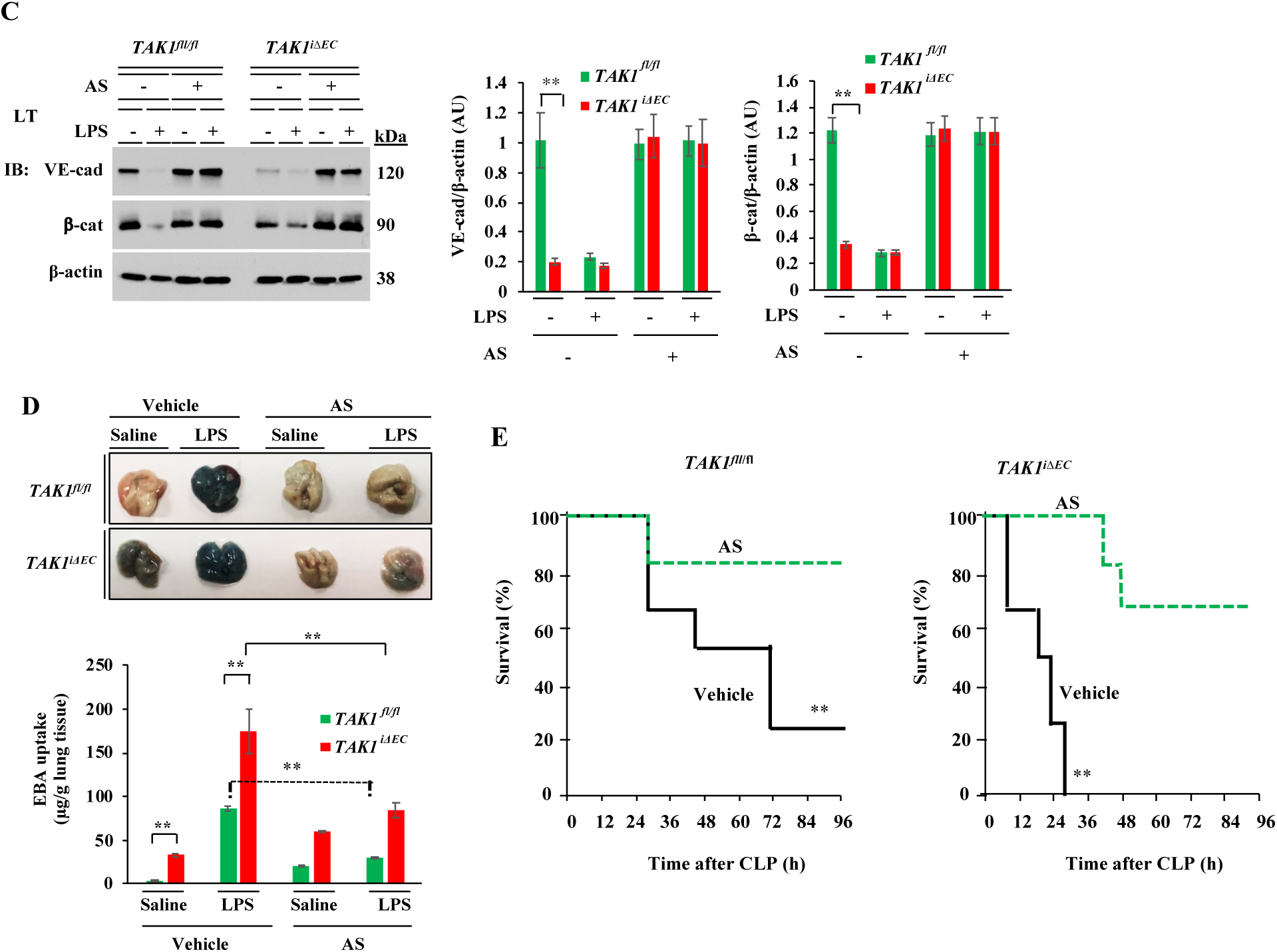
FoxO1 inhibition restores the endothelial barrier and reduces sepsis induced mortality in *TAK1^iΔEC^* mice. **A**, Lung lysates from *TAK1^fl/fl^* and *TAK1^iΔEC^* mice were used for IB to determine expression of AKT and FoxO1. N = 5 mice/genotype; **p<0.001, compared with *TAK1^fl/fl^*mice. **B** and **C**, *TAK1^fl/fl^*and *TAK1^iΔEC^* mice injected i.p. with either vehicle (DMSO) or FoxO1 inhibitor (AS; 5 mg/kg, i.p. one injection per day for 4 consecutive days) were challenged with i.p. LPS (5 mg/kg) or saline for 6 h. Lungs harvested were used for IB analysis. N = 5 mice/genotype/group. **p<0.001, compared with *TAK1^fl/fl^* mice, or *TAK1^iΔEC^* vs *AS+ TAK1^iΔEC^*-LPS. **D,** *TAK1^fl/fl^* and *TAK1^iΔEC^* mice injected i.p. with either vehicle (DMSO) or FoxO1 inhibitor as above were challenged with i.p. LPS (5 mg/kg) or saline for 6 h and used to assess lung vascular leak by measuring EBA uptake. Representative lung images are shown in *top panel*. Quantified results are shown in *bottom panel.* N = 5 mice/genotype/group. **p<0.001, basal *TAK1^fl/fl^ vs TAK1^iΔEC^*; LPS *TAK1^fl/fl^ vs* LPS *TAK1^iΔEC^* mice; LPS *TAK1^fl/fl^ vs TAK1^fl/fl^* + AS; LPS *TAK1^iΔEC^ vs TAK1^iΔEC^*+ AS. **E**, Survival of *TAK1^fl/fl^* and *TAK1^iΔEC^* mice after CLP. Six hour after CLP, mice were injected i.p. with vehicle (DMSO) or AS (FoxO1 inhibitor) (5 mg/kg,i.p) one injection per day for 4 consecutive days. **p<0.001, vehicle injected vs AS injected (log-rank test).

### EC-specific FoxO1 deletion in mice attenuates sepsis-induced lung vascular leak and mortality

To further strengthen the role of TAK1 in regulating endothelial barrier function through FoxO1, we generated EC-restricted *FoxO1* knockout (*FoxO1^ΔEC^*) mice. Here we used *Cre*-dependent Cas9 knock-in mouse model (42) to generate *FoxO1^ΔEC^* mice. First, we generated CRISPR/Cas9-^cdh5-Cre^ mice and then, *FoxO1* was deleted in adult mice ECs by liposome-mediated *in vivo* delivery of the plasmid (pGS) encoding single guide-RNA (sgRNA) to target *m*FoxO1. We used two different sgRNAs (FoxO1-sgRNA-1; FoxO1-sgRNA-2) to target the *m*FoxO1 or a scrambled sgRNA (Sc-sgRNA) as a control. We observed that administration of the plasmid encoding *m*FoxO1-sgRNA-1 effectively disrupted the expression of FoxO1 in lung ECs (Fig. 8A, 8B). Sc-sgRNA or FoxO1-sgRNA-2 had no significant effect on FoxO1 expression. LPS challenge had no effect on FoxO1 expression in FoxO1-sgRNA-1 (*FoxO1^ΔEC^*) mice whereas in WT (Sc-sgRNA-treated) mice, we observed increased expression of FoxO1 (Fig. 8C). Interestingly, LPS challenge caused downregulation of TJ and AJ proteins in WT mice (Fig. 8D, 8E), but LPS had no effect on the expression of TJ and AJ proteins in *FoxO1^iΔEC^* mice (Fig. 8D, 8E). Consistent with these findings, we observed that LPS-induced lung vascular leak (Fig. 8F), and polymicrobial sepsis-induced mortality (Fig. 8G) as markedly reduced in *FoxO1^iΔEC^* mice. These findings collectively suggest that TAK1-mediated FoxO1 inhibition is crucial for endothelial barrier restoration after vascular injury. On the basis of these findings, we propose a model (Fig. 8H) in which TAK1 inhibits the function of GSK3β and FoxO1 via AKT to regulate the endothelial barrier stability.

**Figure 8.**
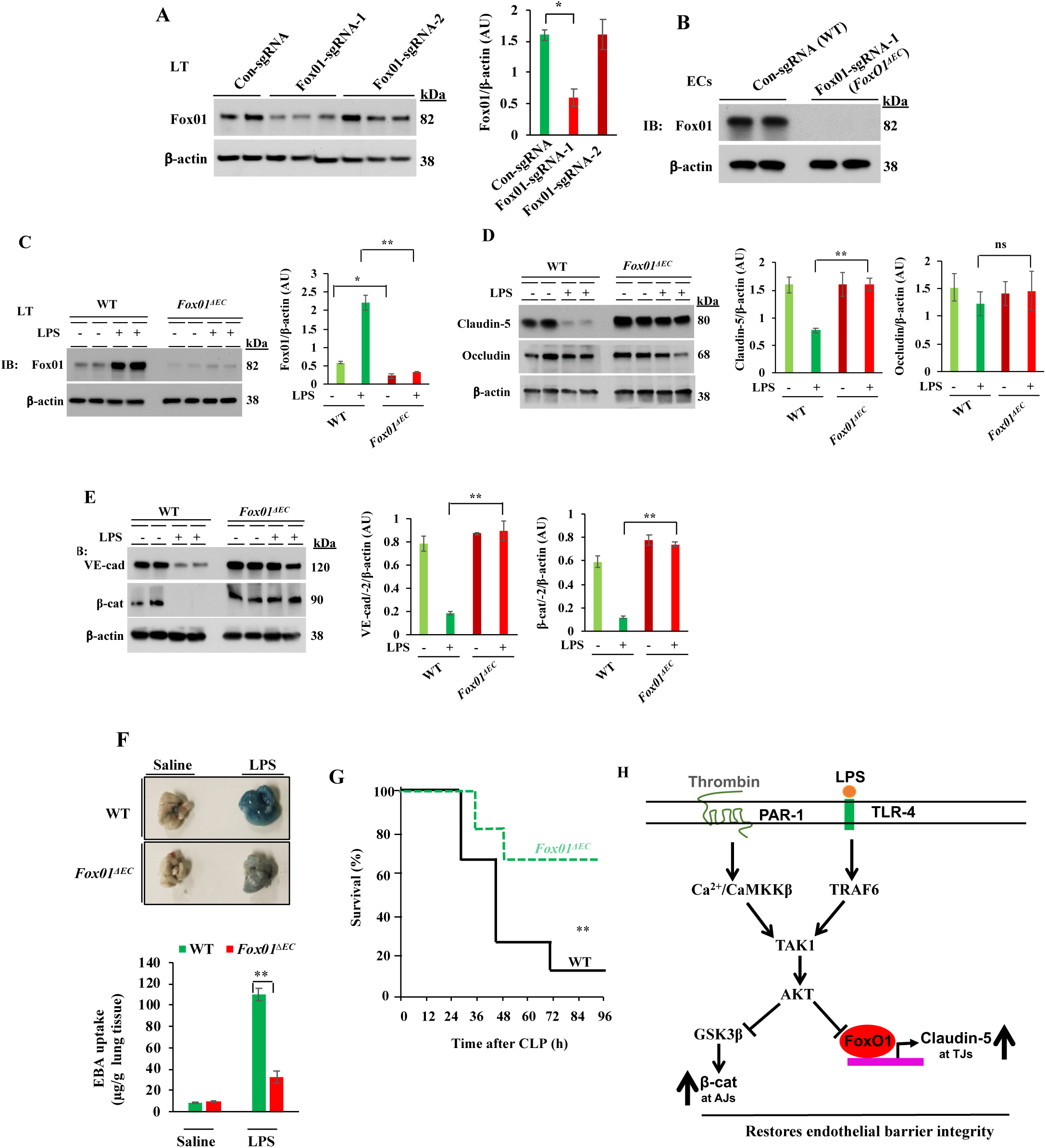
**EC-specific FoxO1 deletion in mice reduces sepsis-induced lung vascular leak and mortality. A**, CRISPR/Cas9-^cdh5-Cre+^ mice were injected with plasmid (pGS-gRNA) encoding sgRNAs (sgRNA-1 or sgRNA-2) to target the *mFoxO1* gene or Control-sgRNA (Scrambled-sgRNA). Lungs harvested were used for IB analysis to determine FoxO1 protein expression. *p<0.01, compared with con-sgRNA. N = 5 mice/group. **B**, Lung ECs from control sgRNA (WT) or FoxO1-sgRNA-1 (*FoxO1^ΔEC^*) mice were used for IB analysis. **C-E**, WT and *FoxO1^ΔEC^* mice were challenged with i.p. LPS (10 mg/kg) or saline for 6 h and then lungs harvested were used for IB analysis to determine expression of FoxO1 (**C**), claudin-5 and occludin (**D**), VE-cad and β-cat (**E**). N= 5 mice/genotype/group. *p<0.01; **p<0.001. **F,** WT and *FoxO1^ΔEC^* mice were challenged with i.p. LPS (10 mg/kg) or saline for 6 h and used for determination of lung vascular leak by measuring EBA uptake. N= 4 mice/genotype/group; **p<0.001. **G,** Survival of WT and *FoxO1^iΔEC^*mice after CLP. N = 8 mice/genotype; **p<0.001, WT vs *FoxO1^Δ^*^EC^ (log-rank test). **H, Model of EC TAK1-dependent mechanisms of repair of endothelial barrier after vascular injury.** In ECs, TAK1 activation downstream of PAR-1 or TLR-4 triggers AKT activation. The activated AKT inhibits GSK3β and transcription factor FoxO1. GSK3β inhibition prevents β-cat degradation, which in turn stabilizes VE-cad at AJs. FoxO1 inhibition increases claudin-5 expression to stabilize TJs.

## Discussion

In this study, we have uncovered the novel functions of TAK1 in regulating vascular homeostasis and repairing the endothelial barrier after vascular injury using our endothelial cell-restricted inducible TAK1 knockout (*TAK1^iΔEC^*) mouse. We observed that *TAK1^iΔEC^*mice exhibited augmented basal lung vascular leak, and that PAR-1 or TLR-4 activation produced uncontrolled lung vascular leak in *TAK1^iΔEC^* mice. Consistent with these results, blocking TAK1 kinase activity caused spontaneous loss of endothelial barrier function due to downregulation of β-cat at AJs in ECs. Importantly, we showed that TAK1 inactivates GSK3β by phosphorylating GSK3β on S9 via AKT activation to restore β-cat at AJs. To provide *in vivo* genetic evidence, we created EC-restricted inducible *GSK3β* knockout (*GSK3β^iΔEC^*) mice and showed that increased β-cat expression in ECs of *GSK3β^iΔEC^* mice, supporting the role of GSK3β in regulating β-cat expression in ECs. Surprisingly, we observed that GSK3β deficiency in ECs induced the formation of β-cat/FoxO1 transcriptional complex, which in turn repressed the expression of the TJ protein claudin-5 to cause endothelial barrier destabilization. Further, we observed augmented expression of FoxO1 while the expression of the FoxO1 negative regulator AKT was suppressed in *TAK1^iΔEC^* mice. Strikingly, EC-restricted FoxO1 deletion of (*FoxO1^ΔEC^*) in mice suppressed sepsis-induced lung vascular leak and mortality. Thus, our results demonstrate for the first time that TAK1 functions as a rheostat in ECs to maintain and repair the endothelial barrier after vascular injury through ATK activation-mediated inhibition of GSK3β and FoxO1 in ECs.

We observed that either inhibition of TAK1 or deletion of TAK1 in ECs, induced loss of β-cat at AJs to cause endothelial barrier destabilization. In support these data, we observed augmented basal as well as thrombin- or LPS-induced phosphorylation and ubiquitination of β-cat in ECs with inhibited TAK1. Consistent with the above findings, thrombin or LPS challenge failed to inactivate GSK3β in TAK1 inhibitor-treated ECs, supporting the concept that TAK1 is essential for GSK3β inhibition-mediated β-cat expression at AJs. Next, we investigated the link between TAK1 and GSK3β in ECs and showed that TAK1 interacts with and activates AKT in response to thrombin or LPS in ECs to phosphorylate S-9 on GSK3β thereby inhibiting GSK3β activity. In further experiments, we showed that, similar to TAK1 inhibition, AKT inhibition also induced loss of endothelial barrier function and augmented thrombin- or LPS-induced responses. Collectively, these results show that TAK1 regulates the endothelial barrier through AKT-dependent inhibition of GSK3β signaling in ECs.

GSK3β is known to participate in many cellular signaling pathways. It regulates the function of several transcription factors critical for inducing inflammation and cytokine production, including NF-*κ*B, AP-1, CREB, NFAT, STAT proteins, and β-cat (43–46). However, GSK3β’s *in vivo* role in ECs is poorly understood. To investigate the *in vivo* role of GSK3β in ECs, we generated tamoxifen inducible EC-restricted *GSK3β* knockout (*GSK3β^iΔEC^*) mice, and unexpectedly we found that GSK3β deficiency in ECs prevented β-cat expression at AJs, which led to defective endothelial barrier function. Knockdown of GSK3β in HLMVECs also mimicked the phenotype of a defective endothelial barrier seen in *GSK3β^iΔEC^*mice. Although β-cat expression was increased in ECs of *GSK3β^iΔEC^*mice, the increased β-cat was primarily bound to transcription factor FoxO1 in the nucleus. Since the β-cat/FoxO1 complex is known to repress the expression of the TJ protein claudin-5 in ECs (5), which induces a defective endothelial barrier (5), we determined the expression of TJ proteins in *GSK3β^iΔEC^* mice and their wildtype counterparts.

Interestingly, we showed that expression of TJ proteins claudin-5 and occludin, and also AJ protein VE-cad were markedly reduced under basal conditions in lungs of *GSK3β^iΔEC^* mice whereas β-cat was upregulated. Next, we observed that FoxO1 inhibitor treatment *in vivo* prevented LPS-induced loss of claudin-5, occludin, VE-cad, and β-cat in WT mice. Importantly, FoxO1 inhibitor treatment, restored the expression of these proteins to a normal level in *GSK3β^iΔEC^* mice and prevented LPS-induced lung vascular leak in both genotypes, thus supporting the hypothesis that the β-cat/FoxO1 complex in ECs causes vascular barrier instability.

FoxO1 expressed in ECs regulates vascular homeostasis (47–51) and its function is tightly regulated by the PI3K-AKT pathway (39–41). AKT phosphorylation of FoxO1 at T-24 expels FoxO1 from nucleus whereas phosphorylation at S-256 inhibits FoxO1 binding to DNA, and thus AKT controls FoxO1 function. The AKT isoform is probably AKT1, the main isoform in ECs that targets FoxO1 (52). We used a pharmacological approach to examine whether TAK1 is required for AKT activation in response to inflammatory stimuli. We showed that TAK1 inhibition blocked both thrombin and LPS induced activation of AKT in ECs. Further, we showed that AKT inhibition blocked thrombin as well as LPS-induced phosphorylation (inactivation) of GSK3β in ECs. Also, we showed that AKT inhibition augmented the thrombin- or LPS-induced endothelial permeability response. These findings are in agreement with the previous report indicating increased vascular permeability in AKT1-null mice (53).

Since we observed loss of endothelial barrier function in *TAK1^iΔEC^* mice, we investigated whether endothelial TAK1 signaling is required for AKT activation during the endothelial repair process, and whether repair of the endothelial barrier involves AKT dependent inactivation of FoxO1 in ECs. We showed that chemical inhibition of TAK1 prevented AKT activation-mediated phosphorylation of FoxO1 at T-24 and S-256. Further, we showed that AKT activated downstream of PAR-1 or TLR4, phosphorylates FoxO1 in ECs. Consistent with these findings, we observed augmented expression of FoxO1 in *TAK1^iΔEC^* mice whereas AKT expression was blocked. To address whether the loss of endothelial barrier function in *TAK1^iΔEC^* mice was due to uncontrolled FoxO1 activation, we inhibited FoxO1 and measured sepsis-induced responses in WT and *TAK1^iΔEC^* mice. We observed that FoxO1 inhibition prevented LPS-induced downregulation of TJ and AJ proteins and lung vascular leak in WT mice. Similarly, FoxO1 inhibition restored TJ and AJ proteins to the basal levels and prevented LPS-induced lung vascular leak in *TAK1^iΔEC^* mice. Importantly, FoxO1 inhibition, markedly reduced CLP-induced mortality in both WT and *TAK1^iΔEC^* mice. Further, we observed that in FoxO1*^ΔEC^* mice, sepsis-induced lung vascular leak and mortality were substantially reduced. These findings collectively support the notion that TAK1 regulates endothelial barrier integrity through AKT-mediated inhibition of GSK3β and FoxO1 in ECs.

TAK1 has been shown to be crucial for mediating inflammation through NF-*κ*B activation in many cell types (21). Recent published studies have demonstrated that TAK1 is also essential to prevent cell death in macrophages, cancer cells, and ECs (27, 28, 32). In our studies, under basal conditions we did not observe endothelial cell death either in TAK1 knockdown ECs or in TAK1-null ECs. We cannot rule out the possibility that during sepsis, TAK1-null ECs undergoes apoptosis to cause vascular barrier disruption. Importantly, chemical inhibition of TAK1 caused spontaneous loss of endothelial barrier function indicating that constitutive activity of TAK1 could regulate endothelial barrier integrity by suppressing the function of GSK3β and FoxO1. Consistent with this notion, we observed that in *TAK1^iΔEC^* mice, the sepsis-induced lung vascular injury response was not reversible, indicating that TAK1-mediated AKT activation during inflammation is critical for repairing the endothelial barrier after vascular injury. In conclusion, we have identified TAK1 as an upstream signaling molecule in ECs that controls the functions of GSK3β and FoxO1 to maintain and repair the endothelial barrier after vascular injury. We believe that this novel pathway controls the restoration rate of endothelial junctions under normal physiological conditions and in the setting of vascular leak. Thus, targeting the TAK1-AKT-GSK3β-FoxO1 axis is a potential therapeutic approach to treat the uncontrolled lung vascular leak seen in sepsis.

## Materials and methods

### Antibodies

Monoclonal antibody (mAb) against TAK1 (cat. no. 4505S), polyclonal antibody (pAb) against p-β-catenin^S33/37/T41^ (cat. no. 9561S), pAb against ubiquitin (cat. no. 3933S), pAb against p-GSK3β^S9^ (cat. no. 9322S), mAb against GSK3β (cat. no. 9315S), mAb against p-AKT^S473^ (cat. no. 4058S), mAb against AKT (cat. no. 4691P), mAb against p-FoxO1^T24/34^ (cat. no. 9464T), mAb against p-FoxO1^S256^(cat. no. 9461T), and mAb against FoxO1 (cat. no. 2880T) were from Cell Signaling Technology (Danvers, MA). pAb against VE-cadherin (cat. no. ab 33168) was from Abcam (Cambridge, MA). pAb against β-catenin (cat. no. sc-7199) and pAb against vWF (cat. no. sc-8068) were obtained from Santa Cruz Biotechnology (Santa Cruz, CA). pAb against occludin (cat. no. 13409-I-AP), and pAb against Lamin B1 (cat. no. 12987-1-AP) were from Proteintech Group Inc (Chicago, IL). pAb against claudin-5 (cat. no. 34-1600) was from Invitrogen and mAb against β-actin (cat. no. A5441) was obtained from Sigma (St. Loius, MO). *In situ* cell death detection (TUNEL Assay) kit, TMR red (Cat no. 12156792910) was obtained from Roche Diagnostic (Indianapolis, IN).

### Other reagents

Protease-activated receptor 1 (PAR-1)-activating peptide (TFLLRNPNDK-NH_2_) was custom synthesized as the C-terminal amide with a purity of > 95% by Genscript (Piscataway, NJ). Scrambled-siRNA (Sc-siRNA) and human (*h*)-specific siRNA to target TAK1, sequences: 5’-GUGCUGACAUGUCUGAAAUTT-3’; 5’-AUUUCAGACAUGUCAGCAC-3’, were custom synthesized by IDT (Coralville, IA). siRNA to target *h*-GSK3β (cat. No. L-003010-00; ON-TARGETplus SMARTpool siRNA sequences: 5’-GAUCAUUUGGUGUGGUAUA-3’; 5’-GCUAGAUCACUGUAACAUA-3’; 5’-GUUCCGAAGUUUAGCCUAU-3’; 5’-GCACCAGAGUUGAUCUUUG-3’) were obtained from Dharmacon (Lafayette, CO). LPS *E. coli* 0111:B4 (cat. no. L3012), TAK1 inhibitor 5Z-7-oxozeaenol (cat. no. 09890), MG-132 (cat. no. 474790), and GSK3β inhibitor SB216763 (cat. no. S3442) were from Sigma (St. Louis, MO). FoxO1 inhibitor AS1842856 (cat. no. 2839) was from Axon Med Chem (Groningen, Netherland). pGS-gRNA plasmid encoding single guide-RNA (sgRNA-1, 5’-CACGGGGGTCAAGCGGTTCA-3’; sgRNA-2, 5’-AATTCGGTCATGCCAGCGTA-3’) to target the *mFoxO1*gene and scrambled sgRNA (Sc-sgRNA) (5’-GCGAGGTATTCGGCTCCGCG-3’) were custom made by Genscript (Piscataway, NJ).

### Methods

#### Mice

*MAP3K7^flox/flox^ (TAK1^fl/fl^)* (33) and *GSK3β^fl/fl^* (54) mice were all described previously. CRISPR/Cas9 knock-in mice (Rosa26-LSL-Cas9 knock-in on B6j [Rosa26-floxed STOP-Cas9 knock-in on B6J]) (42) and B6.Cg-Tg(Cdh5-Cre)7Mlia/J transgenic mice were obtained from Jackson Laboratories (Bar Harbor, ME). All mice were maintained in pathogen-free environment at the University of Illinois Animal Care Facility in accordance with institutional guidelines of the US National Institutes of Health. All animal experiments were performed under the protocol approved by the Institutional Animal Care and Use Committee of the University of Illinois at Chicago.

### Generation of Mutant Mice

#### TAK1^iΔEC^ mice

*TAK1^fl/fl^* mice were bred with *End-SCL-Cre-ER(T)* mice containing tamoxifen-inducible Cre-ER(T) driven by 5’ endothelial enhancer of the stem cell leukemia locus (34, 55) to generate *TAK1^fl/fl-Cre-^* (WT) and TAK1^fl/fl-Cre+^ mice. *TAK1^fl/fl-Cre-^* and TAK1^fl/fl-Cre+^ mice were administered tamoxifen (1 mg/mouse) i.p for 2 consecutive days to generate EC-restricted inducible TAK1 knockout (*TAK1^iΔEC^*) mice.

#### GSK3β ^iΔEC^ mice

A similar protocol described above was used to generate EC-restricted inducible GSK3β knockout (*GSK3β ^iΔEC^*) mice.

#### FoxO1^ΔEC^ mice

We generated CRISPR/Cas9-^cdh5-Cre+^ mice by crossing CRISPR/Cas9 knock-in mice (Rosa26-LSL-Cas9 knock-in on B6j [Rosa26-floxed STOP-Cas9 knock-in on B6J]) (42) with Cdh5-Cre transgenic mice. Next, we generated EC-restricted FoxO1 knockout (*FoxO1^ΔEC^*) mice by liposome-mediated *in vivo* delivery of the pGS-gRNA plasmid encoding sgRNA-1 (5’-CACGGGGGTCAAGCGGTTCA-3’) or sgRNA-2 (5’- AATTCGGTCATGCCAGCGTA-3’) to target the *mFoxO1*gene in CRISPR/Cas9-^cdh5-Cre^ mice. CRISPR/Cas9-^cdh5-Cre^ mice received pGS-gRNA plasmid encoding scrambled sgRNA (Sc-sgRNA) (5’-GCGAGGTATTCGGCTCCGCG-3’) considered as wildtype (42). The liposome and pGS-gRNA plasmid mixture was injected i.v. into CRISPR/Cas9-^cdh5-Cre+^ mice (30 μg of plasmid in 100 μl of liposome suspension/mouse) (56, 57). Four days after sgRNA delivery, mice were used for experiments. Experimental lung injury was induced by systemic LPS (i.p.) in mice (58). Lung injury was assessed by pulmonary capillary filtration coefficient (K_f,c_) measurement using isolated lung preparations (59) and *in vivo* Evans blue dye conjugated albumin (EBA) uptake in lungs (16, 57). Ploymicrobial sepsis was induced by CLP as described (60). Caecum was punctured using 18-gauze needle on 5 different places. For survival studies, mice were monitored four times daily.

### Cells and siRNA transfection

HLMVECs were cultured in EGM-2MV supplemented with 15% FBS as described (16). Mouse (C57BL/6) lung endothelial cells (ECs) were isolated and cultured as described previously (59). Both cell types were used between passages 3 and 6. HLMVECs grown to 70-80% confluence on gelatin-coated culture dishes were transfected with target siRNAs or sc-siRNA as described (16). At 72 h after transfection, cells were used for following experiments. Bone marrow-derived macrophages (BMDMs) from mice were generated by culture of bone marrow cells as described (61). Cell fractionation was performed according to Abcam protocol (https://www.abcam.com/ps/pdf/protocols/subcellular_fractionation.pdf).

### Immunostaining

ECs grown on glass coverslips were washed with HBSS and fixed with 3% paraformaldehyde for 20 minutes. Cells were permeabilized with 0.1% Triton-X-100 for 15 min at 4°C. Next, cells were blocked in 5% goat serum in HBSS for 1 h at room temperature, incubated with primary antibody overnight at 4°C followed by incubation with Alexa-Fluor-conjugated secondary antibody and 4’,6-diamidino-2-phenylindole (DAPI) 1 h at room temperature. In each step, cells were washed three times with HBSS. Coverslips were mounted with prolong gold anti-fade reagent (Invitrogen) on glass slide. Lung tissue sections were immunostained as described (57). Images were acquired with the Zeiss LSM 510 confocal microscope.

### Immunoprecipitation and immunoblotting

ECs grown to confluence treated with or without specific agents were three times washed with phosphate-buffered saline (PBS) at 4^0^C and lysed in lysis buffer (50 mM Tris-HCl, pH7.5, 150 mM NaCl, 1 mM EGTA, 1% Triton X-100, 0.25% sodium deoxycholate, 0.1% SDS, 10 μM orthovanadate, and protease-inhibitor mixture as described (16). Mouse lungs were homogenized in lysis buffer (16). EC lysates or mouse lung homogenates were centrifuged (13,000 X *g* for 10 min) to remove insoluble materials. Clear supernatant collected (300 μg protein) was subjected to immunoprecipitation. Each sample was incubated overnight with 1 μg/ml of the indicated antibody at 4 °C. Next day, Protein A/G beads were added to the sample and incubated at 4 °C for 1 h. Immunoprecipitates were then washed three times with wash buffer (Tris-buffered saline containing 0.05% Triton X-100, 1 mM Na_3_VO_4_, 1 mM NaF, 2 μg/ml leupeptin, 2 μg/ml pepstatin A, 2 μg/ml aprotinin, and 44 μg/ml phenylmethylsulfonyl fluoride) and used for immunoblot analysis. EC lysates, lung tissue homogenates, or immunoprecipitated proteins were resolved by SDS-PAGE on a 4-15% gradient separating gel under reducing conditions and transferred to a Duralose membrane. Membranes were blocked with 5% dry milk in TBST (10mM Tris-HCl pH7.5, 150 nM NaCl, and 0.05% Tween-20) at RT for 1 h. Membranes were then probed with the indicated primary antibody (diluted in blocking buffer) overnight at 4°C. Next, membranes were washed 3x and then incubated with appropriate HRP-conjugated secondary antibody. Protein bands were detected by enhanced chemiluminescence.

### Transendothelial Electrical Resistance Measurement

Real time changes in transendothelial monolayer electrical resistance (TER) was measured to assess endothelial barrier function (62). Confluent endothelial monolayer incubated with 2% FBS containing medium was exposed to indicated inhibitors, thrombin, or LPS. Data are presented as resistance normalized to its starting value zero time.

### Statistical Analysis

Immunoblot bands were quantified using NIH ImageJ software. ANOVA and Student’s *t*-test (2-tailed), and log-rank test were used to determine statistical significance with a *P* value threshold set at <0.05.

## ACKNOWLEDGMENTS

We thank Y.Y. Zhao (University of Illinois at Chicago) and J.R. Woodgett (Samuel Lunenfeld Research Institute, Toronto) for *GSK3β^fl/fl^* mice. This work was supported by National Institutes of Health Grants R01 GM-117028, R01 HL-128359, and R01 HL-122157.

## AUTHOR CONTRIBUTIONS

SCR, DS, DMW, ABM, and CT designed the research. CSR, DS, DMW, SMV, performed experiments. CT, CSR, DS, DMW, SMV analyzed and interpreted the results. DS, CSR, ABM, and CT wrote the manuscript. CT approved the manuscript.

