## Supplemental Data for "TAK1 Regulates Endothelial Integrity Through Stabilization of Junctions via GSK3β and FoxO1"

Sushil C. Regmi, Dheeraj Soni, Dong-Mei Wang, Stephen M. Vogel, Asrar B. Malik, and Chinnaswamy Tiruppathi*
Department of Pharmacology and Center for Lung and Vascular Biology,
College of Medicine, University of Illinois, Chicago, IL, USA

**Running title:** TAK1 inhibits GSK3β and FoxO1

**SCR and DS:** These authors contributed equally.

**Correspondence to:** Chinnaswamy Tiruppathi, Ph.D., Department of Pharmacology (M/C868), College of Medicine, University of Illinois, 835 South Wolcott Ave, Chicago, IL 60612; Telephone: 312-355-0249; Fax: 312-996-1225;

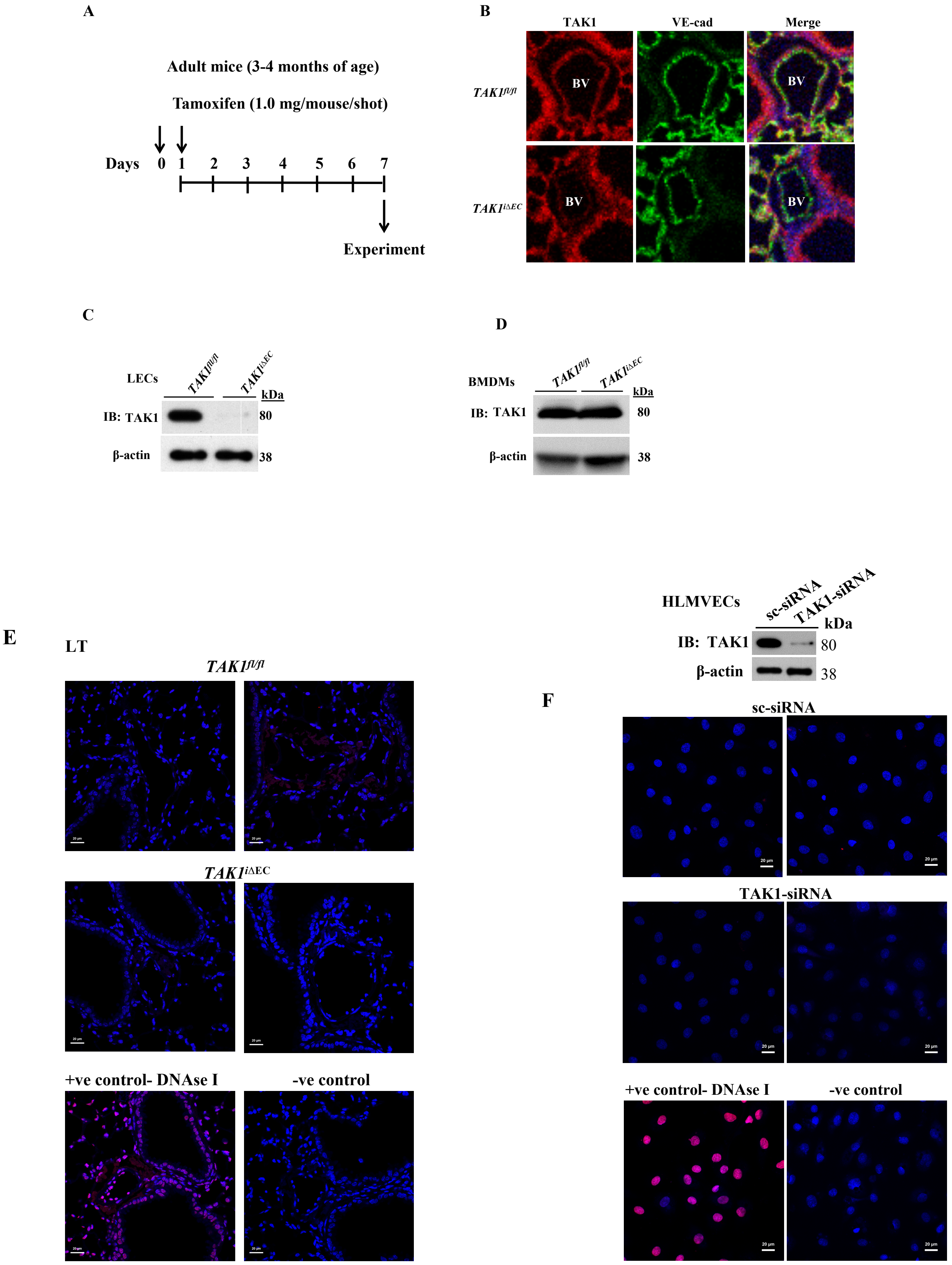
A
Adult mice (3-4 months of age)
Tamoxifen (1.0 mg/mouse/shot)
Days 0 1 2 3 4 5 6 7
Experiment
B
TAK1
VE-cad
Merge
TAK1fl/fl
TAK1iΔEC
BV
C
LECs
TAK1fl/fl
TAK1iΔEC
kDa
IB: TAK1
80
β-actin
38
D
BMDMs
TAK1fl/fl
TAK1iΔEC
kDa
IB: TAK1
80
β-actin
38
HLMVECs
sc-siRNA
TAK1-siRNA
kDa
IB: TAK1
80
β-actin
38
E
LT
TAK1fl/fl
TAK1iΔEC
+ve control- DNAse I
-ve control
F
sc-siRNA
TAK1-siRNA
+ve control- DNAse I
-ve control

**Supplemental Figure 1. EC-restricted tamoxifen-inducible TAK1 deletion in mice.**

**A,** Shows tamoxifen administration schedule. *TAK1*$^{fl/fl}$ and TAK1$^{fl/fl/Scl\text{-}CreERT2+}$ mice were i.p. injected to delete TAK1 in TAK1$^{fl/fl/Scl\text{-}CreERT2+}$ mice (*TAK1*$^{i\Delta EC}$).

**B**, Lung sections prepared from *TAK1*$^{fl/fl}$ and *TAK1*$^{i\Delta EC}$ mice were stained with antibodies specific to TAK1 and VE-cad. DAPI (blue). BV, blood vessel.

**C**, Lung endothelial cells (LECs) from *TAK1*$^{fl/fl}$ and *TAK1*$^{i\Delta EC}$ mice were used for IB analysis to determine TAK1 expression.

**D,** Bone marrow-derived macrophages (BMDMs) from *TAK1*$^{fl/fl}$ and *TAK1*$^{i\Delta EC}$ mice were used for IB analysis to determine TAK1 expression.

**E**, Lung sections from *TAK1*$^{fl/fl}$ and *TAK1*$^{i\Delta EC}$ mice were used for TUNEL staining. Images were analyzed by confocal microscopy.

**F**, HLMVECs transfected with sc-siRNA or TAK1-siRNA were used for TUNEL staining.

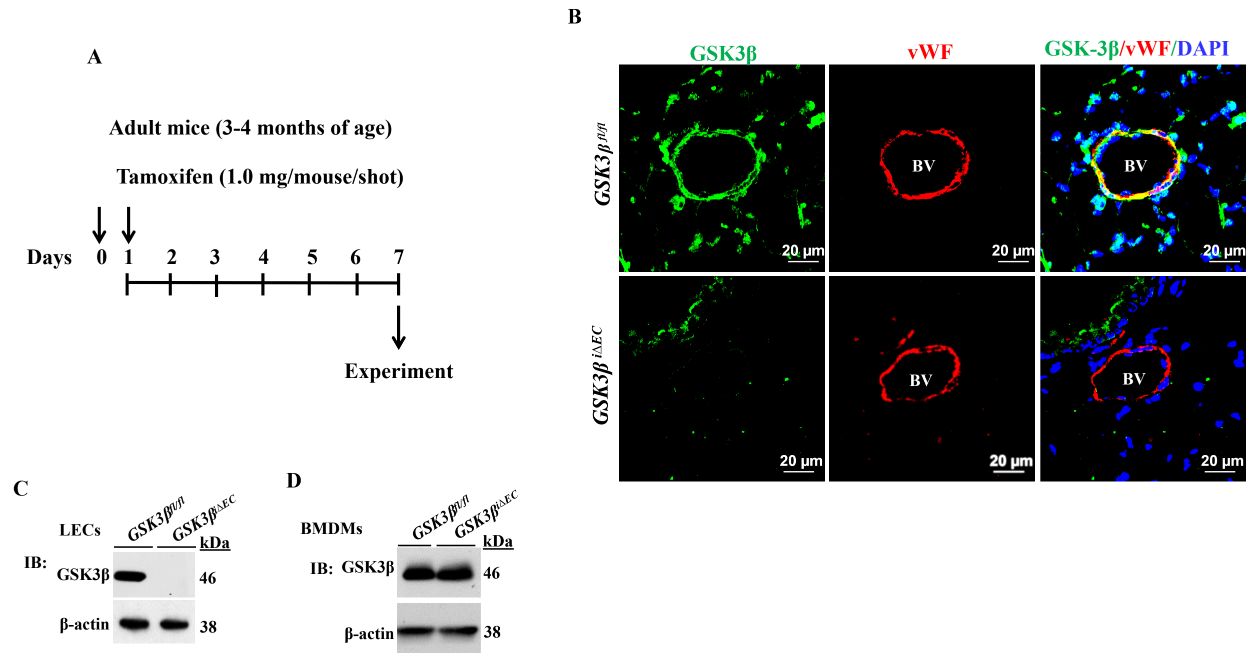


**Supplemental Figure 2. EC-restricted tamoxifen-inducible GSK3β deletion in mice.**

**A,** Shows tamoxifen administration schedule. *GSK3β*$^{fl/fl}$ and *GSK3β*$^{fl/fl/Scl\text{-}CreERT2+}$ mice were i.p. injected to delete GSK3β in *GSK3β*$^{fl/fl/Scl\text{-}CreERT2+}$ mice (*GSK3β*$^{i\Delta EC}$).

**B**, Lung sections prepared from *GSK3β*$^{fl/fl}$ and *GSK3β*$^{i\Delta EC}$ mice were stained with antibodies specific to GSK3β and vWF (EC-marker). DAPI (blue). BV, blood vessel.

**C,** Lung endothelial cells (LECs) from *GSK3β*$^{fl/fl}$ and *GSK3β*$^{i\Delta EC}$ mice were used for IB analysis to determine GSK3β expression.

**D,** Bone marrow-derived macrophages (BMDMs) from *GSK3β*$^{fl/fl}$ and *GSK3β*$^{i\Delta EC}$ mice were used for IB analysis to determine GSK3β expression.
